# Coexistence of many species under a random competition-colonization trade-off

**DOI:** 10.1101/2023.03.23.533867

**Authors:** Zachary R. Miller, Maxime Clenet, Katja Della Libera, François Massol, Stefano Allesina

**Author notes:** These authors contributed equally to the work (Zachary R. Miller, Maxime Clenet). These authors jointly supervised the work (François Massol, Stefano Allesina).

## Abstract

The competition-colonization trade-off is a well-studied coexistence mechanism for metacommunities. In this setting, it is believed that coexistence of all species requires their traits to satisfy restrictive conditions limiting their similarity. To investigate whether diverse metacommunities can assemble in a competition-colonization trade-off model, we study their assembly from a probabilistic perspective. From a pool of species with parameters (corresponding to traits) sampled at random, we compute the probability that any number of species coexist and characterize the set of species that emerges through assembly. Remarkably, almost exactly half of the species in a large pool typically coexist, with no saturation as the size of the pool grows, and with little dependence on the underlying distribution of traits. Through a mix of analytical results and simulations, we show that this unlimited niche packing emerges as assembly actively moves communities toward overdispersed configurations in niche space. Our findings also apply to a realistic assembly scenario where species invade one-at-a-time from a fixed regional pool. When diversity arises de novo in the metacommunity, richness still grows without bound, but more slowly. Together, our results suggest that the competition-colonization trade-off can support the robust emergence of diverse communities, even when coexistence of the full species pool is exceedingly unlikely.

## Introduction

All organisms face physical and evolutionary constraints that limit simultaneous optimization of different fitness components^1^. These constraints create trade-offs, which can foster coexistence by making multiple distinct trait combinations ecologically viable^2,3^. A number of trade-offs are thought to contribute to maintaining the incredible diversity of natural communities^4–9^. Perhaps the best-known of these is the competition-colonization (CC) trade-off, wherein dominant competitors are constrained to be relatively poor colonizers of available habitat, due to low fecundity, dispersal, or growth rates^10–12^. Weaker competitors may then persist by rapidly reaching and reproducing in open habitat patches, staying one step ahead of competitive exclusion at the landscape scale^3,13^.

The essential dynamics of this scenario can be captured in a simple patch-occupancy model^14–16^. Tilman^17^ showed that this minimal model can potentially explain the coexistence of arbitrarily many species along a single trade-off axis. Independently, May and Nowak^18,19^ obtained similar results, using an identical mathematical model, in an epidemiological context. These results are somewhat surprising, because the model assumes a strict competitive hierarchy; in a spatially well-mixed setting, only the single best competitor would persist.

Despite this theoretical proof of concept, the extent to which the CC trade-off can explain natural biodiversity remains murky. Empirical tests have yielded mixed results^13,20–23^, although evidence for this trade-off – and its role in maintaining coexistence – has been found in many taxa, including plants^5,24–26^, bacteria^27,28^, fungi^29,30^, birds^31^, insects^32^, and taxonomically mixed communities^33,34^. From a theoretical perspective, there has been significant debate over the specific assumptions underlying CC trade-off models, and how these assumptions affect predicted coexistence^12,13,35,36^.

An even more fundamental question, though, is whether diverse communities can emerge and coexist under the CC trade-off with realistic limits to colonization rates and no fine-tuning of species traits (model parameters)^37^. In the basic patch-occupancy model developed by Tilman, each species excludes inferior competitors within a range of colonization rates similar to its own^17,18^. This limiting similarity suggests that the trade-off axis could quickly become saturated, especially when there is an upper limit to species’ colonization abilities^17^. However, the degree of limiting similarity is an emergent property of this model, making it difficult to predict whether or how niche space saturates. Later work showed that arbitrarily many coexisting species can in fact be packed into a finite range of colonization rates, but this construction requires precisely spaced colonization rates^38^. Small perturbations to such a fine-tuned packing may cause the loss of many species. Indeed, from a probabilistic viewpoint, the likelihood that a given set of species will coexist decreases exponentially with its size^12,39^, reminiscent of the classic complexity-stability paradox^40,41^.

To clarify the typical behavior of the CC trade-off, we adopt a community assembly perspective. Rather than asking whether a particular set of species can coexist, we examine metacommunities that emerge through the model dynamics from an initial random pool^41–43^. This probabilistic approach is a powerful way to identify characteristic outcomes in complex models^44–46^. We investigate not only the expected number of coexisting species from a pool of *n* species, but also the distribution of this quantity, as well as typical features of self-assembled metacommunities. We find that on average half of the species coexist, and these persistent communities become overdispersed in niche space. These properties show remarkably universal behaviors for a range of possible trait distributions in the species pool. Finally, we consider how these results might change when metacommunities are assembled one-species-at-a-time from a persistent regional pool or through the sequential invasion of unique species.

## Results

We focus on the well-known CC trade-off model^17^

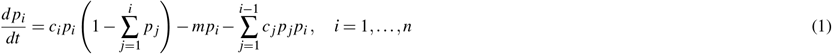

describing the fraction of patches (*p*_*i*_ ≥ 0) occupied by each of *n* species in a metacommunity (∑_*i*_ *p*_*i*_ ≤1). Here, *m >* 0 is the local extinction rate, which we assume to be equal for all species, and *c*_*i*_ *>* 0 is the colonization rate of species *i*. We study the behavior of this model when colonization rates are independent samples from a fixed distribution, ordered such that *c*_*i*_ *< c* _*j*_ for all *i < j*. This distribution defines how colonization ability tends to increase with decreasing competitive rank. Given a pool of species defined by their colonization rates, we then ask which species coexist through the dynamics (Fig. 1).

**Figure 1.**
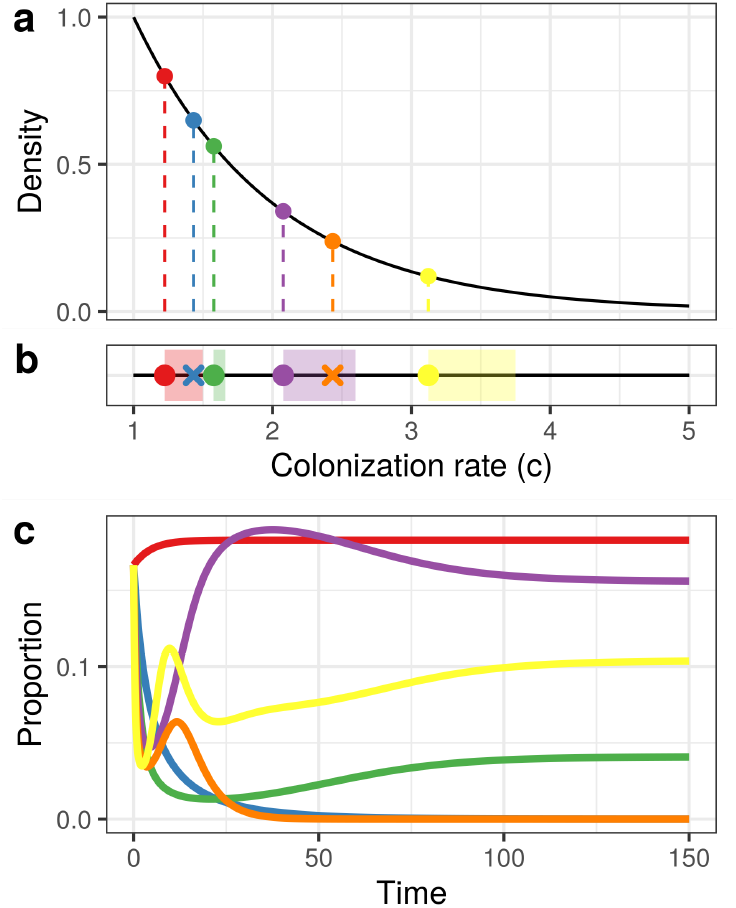
Example species pool and metacommunity dynamics. (a) A set of six species with colonization rates sampled independently from a fixed distribution (here, an exponential distribution with rate 1, truncated at *m* = 1). (b) Proceeding from lowest to highest colonization rate (equivalently, highest to lowest competitive ability), each species excludes inferior competitors within a “niche shadow” (shaded regions). The length of each niche shadow depends on the distance from the species casting it to the previous niche shadow. Species that fall within the niche shadow of a superior competitor will be excluded (indicated by an X). (c) Consistent with this prediction, the blue and orange species go extinct in a simulation of the model dynamics, while the other four stably coexist.

First, we consider the mechanics of coexistence. Every non-negative equilibrium is stable in this model, and thus coexistence is determined solely by which species are excluded (i.e. driven to zero occupancy at equilibrium) by superior competitors^17,39^. Proceeding down the competitive hierarchy, each species *i* excludes inferior competitors with colonization rates below a threshold, *ℓ*_*i*_ (Fig. 1b). The length of this exclusion interval (*c*_*i*_, *ℓ*_*i*_), which is called the “niche shadow”, depends recursively on the preceding one – as *c*_*i*_ approaches its own lower bound (that is, *ℓ*_*i*−1_), the shadow cast by species *i* shrinks^12,38^. In ecological terms, when species *i* has higher niche overlap with superior competitors, it is competitively suppressed, and less able to suppress its own inferior competitors. As a result, its niche shadow shrinks, providing more opportunity for coexistence of species that are slightly worse competitors but better colonizers. Thus, as the species pool grows, there are two opposing forces that shape the potential for coexistence: (i) each species adds a new niche shadow along the trade-off axis, reducing space for coexistence, but (ii) as colonization rates become more densely packed, the typical niche shadow shrinks, increasing space for coexistence^38^.

Surprisingly, we find that these two forces tend to balance out precisely. For a wide range of colonization rate distributions, we observe that niche shadows shrink as 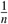, so that each coexisting species excludes one competitor on average. As a result, the number of species that persist in the assembled metacommunity, which we denote by *S*, is typically half of the initial pool. Fig. 2 shows this proportional scaling for species pools ranging from two to more than one thousand species. We present results for four possible distributions of colonization rates – Uniform, Exponential, Pareto (power-law), and Triangular (see SI Sections 2 and 6 for details and alternative distributions). We select these distributions to highlight scenarios with very different qualitative features, such as the presence or absence of a maximal colonization rate, and left- or right-skew. Notably, the number of coexisting species scales as 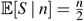 *without saturation* in all cases, regardless of whether the distribution of colonization rates is unbounded (Exponential and Pareto distributions) or bounded (Uniform and Triangular distributions).

**Figure 2.**
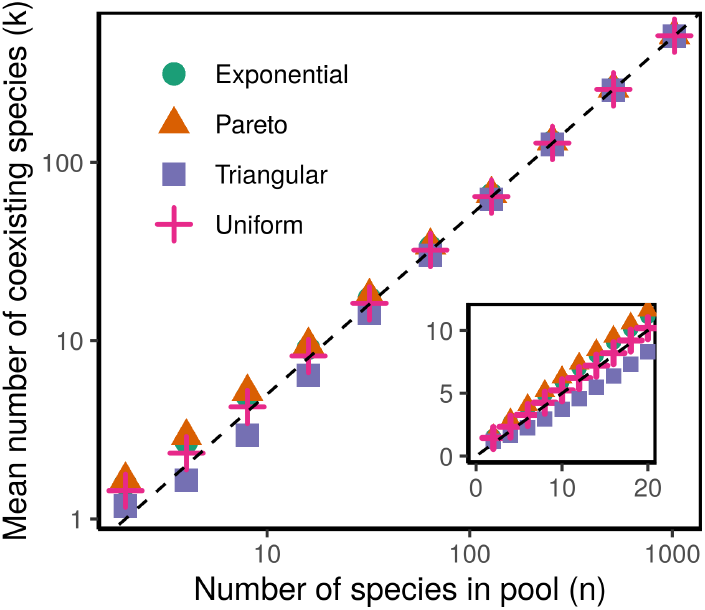
The mean number of coexisting species, 𝔼[*S*| *n*], from a random pool of *n* species grows as 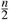 for different distributions of colonization rates. For large *n*, all four distributions converge; for small *n*, 𝔼[*S n*] scales approximately linearly with *n*, but different distributions fall slightly above or below the 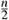 prediction (dashed line), depending on the tail behavior of the distribution. Note the log-log scale of the main figure; the inset highlights small *n* results on a standard scale. Points are averages over 10^5^ random realizations for each distribution and value of *n*.

To better understand this lack of saturation, we examine the full distribution of the number of persisting species. The probability that exactly *s* species coexist from a pool of size *n*, denoted *P*(*S* = *s*| *n*), can be approximated rigorously for the Uniform and Exponential distributions, and heuristically for a much broader class of distributions. We find that, as *n* becomes large, *P*(*S* = *s* | *n*) becomes very close to the binomial distribution 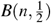. More precisely, because the best competitor always persists in the final metacommunity, the number of persistent species among the remaining *n* 1 is distributed as 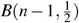. Fig. 3 shows this convergence for all four example distributions. The binomial distribution is tightly peaked around its mean.

**Figure 3.**
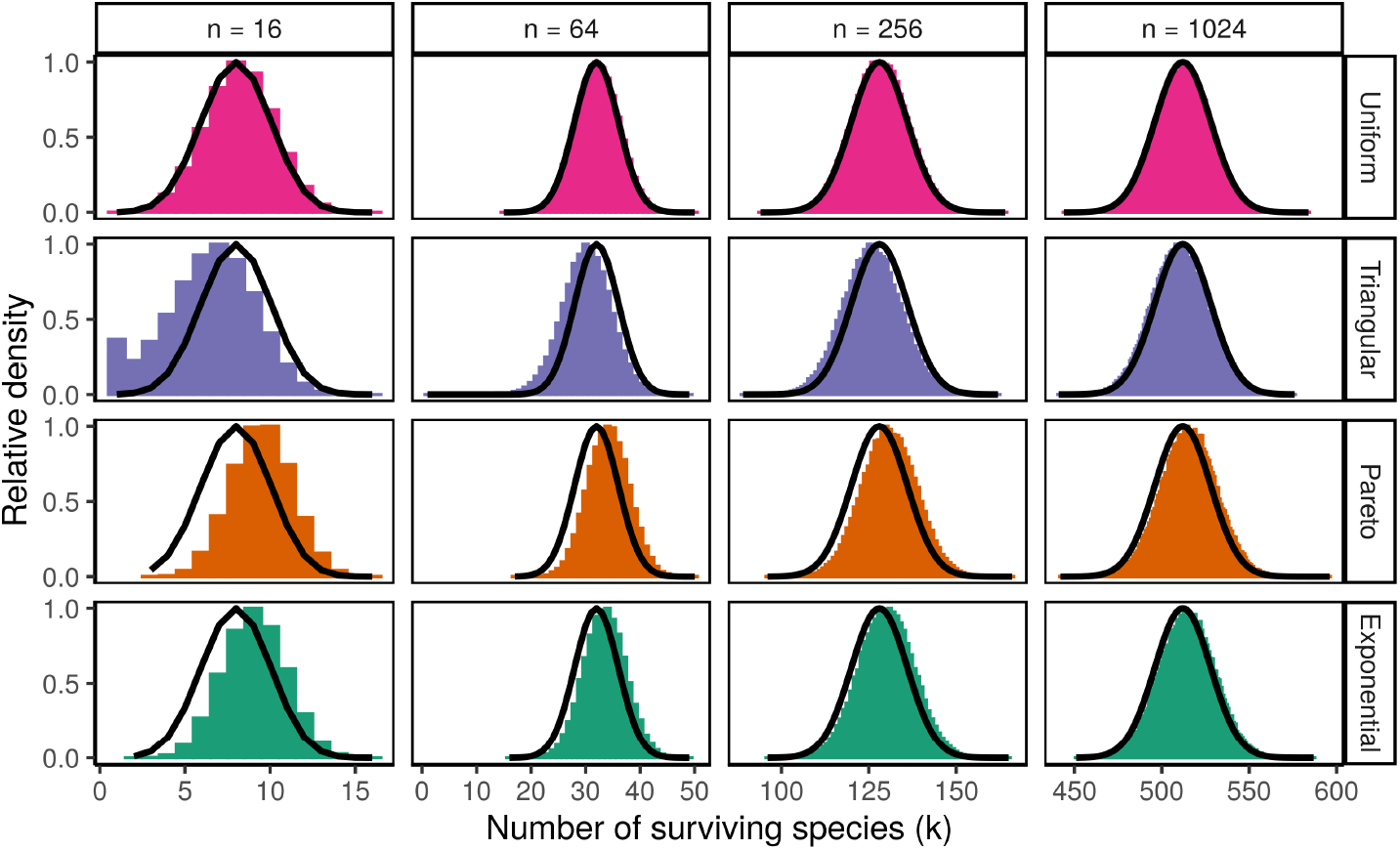
Distributions of the number of persistent species, *S*, from a random pool of *n* species. For large *n, P*(*S* = *s*| *n*) is well approximated by the binomial distribution 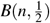 (black curve) regardless of how colonization rates are distributed in the pool. *P*(*S* = *s*| *n*) is nearly binomial even for moderate *n*, and converges very rapidly for the Uniform distribution. Histograms represent the outcomes of 10^5^ random realizations for each distribution and value of *n*. Note that the maximum density in each panel is rescaled to 1 to facilitate comparison across different values of *n*.

This implies that coexistence of all *n* species is extremely unlikely, in agreement with existing theoretical predictions^12,39^ – but so too is the exclusion of many species. Instead, the number of persistent species is nearly always about one half of the pool. Although our analytical results are asymptotic in nature, we find that the binomial distribution provides a good approximation even for moderate *n*. Additionally, Fig. 2 (inset) shows that the mean number of coexisting species already grows linearly (and with slope near 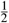) for *n <* 20. In SI Section 2, we show that deviations from the predicted binomial behavior at small *n* (e.g. in Fig. 3 for *n* = 16) can be explained by specific features of the colonization rate distributions. In particular, notice that the Triangular distribution, which has low density near *m*, shows a bias toward fewer coexisting species, while the Exponential and Pareto distributions, which have long upper tails, show slightly increased coexistence. In general, we can show that excess competitive exclusion arises in any lower tail of the colonization rate distribution, while excess coexistence arises in any upper tails, although these deviations typically have negligible effects on the overall metacommunity richness for large *n*.

The appearance of the binomial distribution might suggest that each species in the pool (aside from the best competitor) persists independently with probability 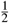, as if we flipped a fair coin to determine the persistence of each species. Consistent with this intuition, we find that the marginal distribution of colonization rates in each assembled metacommunity is identical to the distribution in the corresponding pool (Fig. 4a). In other words, the ecological dynamics induce no significant bias toward the persistence of better competitors or colonizers. However, for all distributions, the *spacing* of coexisting species along the CC trade-off axis shows a marked and consistent signature of the dynamical pruning (Fig. 4b). Compared to a random subset of species from the pool, the spacing between consecutive persisting species (i.e. *c*_*i*_ −*c*_*i*−1_ in the coexisting metacommunity) is substantially more even, with fewer small or large gaps.

**Figure 4.**
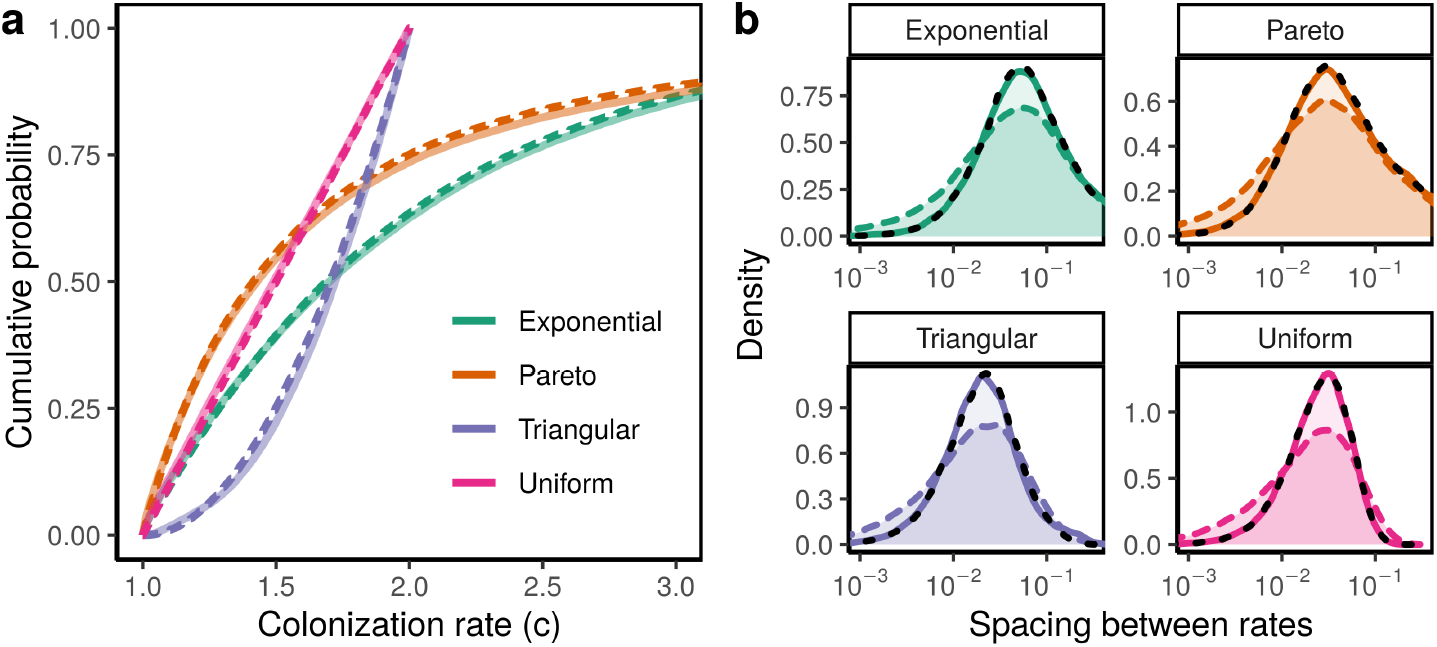
Comparing the distribution of colonization rates in assembled metacommunities vs. the random pool. (a) The marginal distribution of colonization rates in assembled metacommunities (solid lines) closely matches the distribution in the pool (dashed lines). For each distribution, we plot the empirical distribution function of colonization rates for 10^3^ random pools of *n* = 64 species. (b) The distribution of spacings between consecutive colonization rates is more peaked for the assembled metacommunities (solid lines) than for a random set of species of the same size from the corresponding pools (colored dashed lines). In particular, assembled metacommunities are less likely to have very small spacings, compared to a random sample from the pool. In each case, the realized spacings closely match the distribution of spacings between every other species in the pool (black dashed lines). Note that spacings are shown on a log scale to highlight behavior at small values.

For Uniformly distributed pools, we can compute this distribution of niche spacings analytically, and indeed it differs qualitatively from the naive expectation of independent sampling from the pool (see SI Section 3). Under independent random sampling, the theoretical distribution of spacings has a mode at zero, while the realized spacings in the coexisting metacommunity have a non-zero mode; in fact, this distribution has no density at all at zero, indicating substantial repulsion between the assembled colonization rates. Furthermore, we find that the distribution of spacings in the final metacommunity is identical to the distribution of spacings between *every other* colonization rate (i.e. *c*_*i*_ − *c*_*i*−2_) in the species pool – the most evenly-spaced way to choose 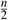 species from the pool on average. Based on this analytical result, we conjecture that this is the general behavior for any distribution of colonization rates. In Fig. 4b, we show that our conjecture matches the simulation results nearly perfectly. Thus, while assembly does not distort the marginal distribution of colonization rates, it does induce correlations between species at small scales along the trade-off axis, moving the assembled metacommunity toward more evenly-spaced configurations, which make coexistence possible^38^.

All of our results so far are obtained under the assumption that the full species pool is introduced simultaneously in the landscape. While this kind of assumption is typical for theoretical studies of coexistence^17,41,44,45^, it is unlikely to reflect community assembly in most natural systems^43,47^. This motivates us to consider whether the ecosystem dynamics behave differently under more realistic assembly scenarios, where the metacommunity is assembled through sequential species invasions^42,48^. At one extreme, we consider a case where species enter the local landscape one-at-a-time from a fixed regional pool, with re-invasion of the same species possible at different points in time. In more mathematical terms, this corresponds to sampling with replacement from a finite species pool. In this scenario, we can prove that the metacommunity will eventually reach a non-invasible composition identical to the coexisting metacommunity under our previous all-at-once assembly assumption^47^ (see SI Section 4.1 for details). For example, in Fig. 5a, we show that when species from the pool illustrated in Fig. 1 are introduced one-at-a-time – rather than all at once – the final outcome is the same (compare Fig. 1c and Fig. 5a after *t* = 1000). This implies that, given sufficient time for assembly to occur, all of our results linking the regional pool and local metacommunity will apply.

**Figure 5.**
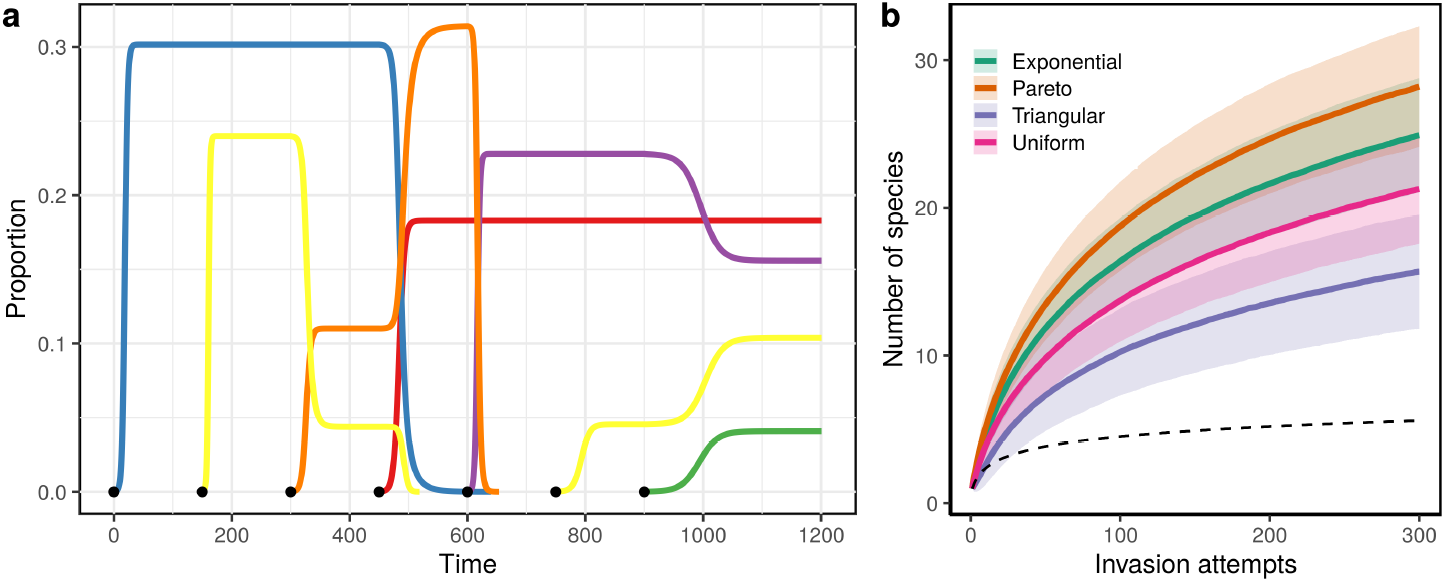
Two different one-at-a-time assembly scenarios. (a) When species invade one-at-a-time from a fixed pool, the metacommunity eventually converges on the same non-invasible equilibrium as if all species were introduced simultaneously. Here, one species is chosen to invade every 150 time units (black dots) from the pool in Fig. 1. As in Fig. 1, the blue and orange species eventually go extinct, and the four others coexist at the same equilibrium occupancies as in Fig. 1c. Note that species present in the final metacommunity may be transiently excluded during the assembly process, only to re-invade later (e.g. yellow species). (b) Accumulation of species richness in the de novo invasion scenario. Colored lines show the mean number of coexisting species as a function of the number of invasion attempts. Shading indicates *±*1 standard deviation. Mean diversity accumulates faster than a logarithmic lower bound (dashed line), but asymptotically these curves grow logarithmically (see SI Section 4.2). Curves represent statistical summaries of 10^5^ random assembly trajectories.

In another limiting case, new species enter the ecosystem through de novo invasion. We assume that invasion events are rare, so that the metacommunity dynamics reach equilibrium between each invasion, and in this scenario each new species is sampled independently from a fixed, underlying distribution of colonization rates (i.e. sampling from an infinite pool). A similar scenario was previously studied by Nowak and May in the context of pathogen evolution^18,19^. In agreement with their analysis, we find that the expected number of coexisting species increases logarithmically with the number of invasion events^18^.

This richness-accumulation relationship is shown in Fig. 5b for different colonization rate distributions. To more rigorously characterize this relationship, we derive the lower bound log(*τ/*2 + 1) for the mean richness after *τ* invasion attempts, regardless of the distribution of colonization rates (see SI Section 4.2). Our simulations substantially exceed this bound, but grow at the same pace asymptotically.

This unbounded increase in diversity is qualitatively consistent with our central finding that the CC trade-off does not saturate. However, as noted by Nowak and May, the logarithmic growth can be extremely slow once diversity is high, making it difficult to distinguish from saturation in practice. This assembly scenario behaves somewhat differently from our previous cases because the strong hierarchical interactions in this model create historical contingency. While there are no true priority effects under this model (i.e., there is a unique equilibrium corresponding to any fixed set of species, regardless of their order of arrival), whether a particular invader can enter the community depends on which set of species has previously invaded. We assume that unsuccessful invaders decline to extinction and never re-invade, so many species are lost from the metacommunity through the course of assembly.

Interestingly, in the de novo invasion scenario, the distribution of colonization rates has a substantial effect on the rate of diversity accumulation. Fig. 5b shows that the absolute difference in mean richness between different distributions grows with the number of invasion attempts (see also Fig. S8). While the shape and asymptotic behavior of these curves is similar, after 300 invasion attempts metacommunities with Pareto-distributed colonization rates have nearly twice as many species as Triangular-distributed metacommunities, on average. As we found for the convergence of *P*(*S* = *s* | *n*) in the all-at-once assembly scenario, this difference arises from the tail behaviors of the colonization rate distributions. Distributions with long upper tails (here the Pareto and Exponential) support faster accumulation of species through de novo invasion than do the distributions with lower tails (Triangular) and bounded support (Uniform).

## Discussion

We asked whether species-rich metacommunities can emerge and coexist under the CC trade-off through ecological assembly. Despite intuitive arguments for the saturation of the trade-off axis^12,17^, we showed that arbitrarily large sets of species can coexist through the model dynamics. No fine-tuning is necessary to produce highly diverse metacommunities, given a large enough species pool. These results hold for a wide range of colonization rate distributions, including cases where there is an upper limit to colonization ability. In these cases, the trade-off axis simply becomes more and more finely partitioned as the metacommunity grows.

Remarkably, species richness at the metacommunity scale is not only generically non-saturating, but also exhibits nearly universal properties when diversity is high (large *n*). For quite different colonization rate distributions, the size of the coexisting metacommunity becomes binomially distributed, implying that in nearly all cases a constant fraction (near one half) of species will coexist. Our numerical simulations highlight that this behavior becomes typical for ecologically relevant pool sizes on the order of tens of species.

We also showed that assembly shapes coexisting metacommunities in consistent ways. These effects are somewhat subtle, highlighting the value of theoretical models for clarifying the signatures of assembly processes in nature. For example, the marginal distribution of colonization rates in the assembled metacommunities is indistinguishable from the distribution in the pool, an apparently “neutral” pattern^49,50^. However, colonization rates in the coexisting metacommunities show fine-scale repulsion, a clear statistical fingerprint of community assembly^42,50,51^. Notably, for Uniformly distributed colonization rates, this repulsion phenomenon can be characterized analytically, providing a uniquely tractable model for the self-organization of assembling communities in niche space. In SI Section 5, we examine another interesting contrast by comparing probability of persistence and average occupancy in the coexisting metacommunity, both as a function of competitive rank. Except for small deviations at the upper and lower extremes of the competitive hierarchy, all species are equally likely to be found in the coexisting metacommunity; however, when stronger competitors persist, they tend to have higher occupancy^38^. Moreover, the species occupancy distribution in the coexisting metacommunity strongly depends on the distribution of colonization rates, while the probability of persistence does not. Our probabilistic approach is instrumental for dissecting these unintuitive patterns. Our basic conclusions are robust across distinct assembly scenarios, although we find that diversity accumulates much more slowly when species cannot re-invade. Further exploration of detailed one-at-a-time assembly processes remains an important avenue for connecting our theoretical results to natural systems, where multiple timescales and sources of species invasions will usually be at play. However, the overarching result that niche packing does not fundamentally limit coexistence in this model suggests it may be important to consider the mechanisms that supply diversity (evolution, immigration, and regional pools) in order to understand what sets observed levels of biodiversity^37,42,43,52,53^.

It will also be important to ask whether these results hold, at least qualitatively, for more general versions of the trade-off model. The basic CC trade-off model that we consider has been critiqued for relying on overly simplistic assumptions^35,36^. In particular, this model assumes that worse colonizers always out-compete better ones, regardless of how similar their colonization rates are (*perfectly asymmetric competition*); that propagules of superior competitors can displace inferior competitors (*displacement competition*); and that there is no spatial structure linking patches (*global dispersal*). Calcagno *et al*.^12^ studied a more complex CC trade-off model relaxing the first two assumptions, and found that the effects of these changes were nuanced, but did not fundamentally alter the capacity of the trade-off to maintain coexistence. Moreover, (nearly) perfectly asymmetric competition is not unknown in the empirical literature^36,54,55^. However, the assumption of asymmetric competition does constrain the possible dynamics, for example precluding the emergence of “nearly neutral” trait clusters, as found in other theoretical studies of assembly^56^. Regarding displacement competition, Calcagno *et al*. showed that relaxing this assumption can actually favor coexistence. This outcome is especially likely when competition is not perfectly asymmetric, suggesting that these assumptions might “cancel out” to some extent. And while considering explicit spatial structure would undoubtedly affect our quantitative results, theory generally indicates that this would only increase the potential for coexistence, strengthening our qualitative conclusions^57^.

Although our analysis reveals outcomes that are quite robust to different colonization rate distributions (and different qualitative features such as the presence or absence of a maximum colonization rate), it does rely fundamentally on the existence of a strict CC trade-off. Whether this is generally the case in natural metacommunities is a question that can only be answered by systematic empirical study. However, our results suggest that when a strong trade-off is present, coexistence of many species is not only possible but typical. Some authors have suggested that other trade-offs, such as competition-defense or colonization-persistence tradeoffs, are more likely to influence coexistence in natural ecosystems^3,8,9,36,58^. It would be interesting to apply a probabilistic assembly perspective, as we showcase here, to study the typical behavior of other types of trade-offs. We hypothesize that trade-offs between other trait combinations, by equalizing fitness across a trait axis, may similarly permit non-saturating niche packing. In fact, a recent theoretical analysis found strikingly similar outcomes in forest dynamics models featuring trade-offs in competition for light^59^.

This last, admittedly speculative, suggestion is bolstered not only by the analysis of other trade-off models, but also by the surprising similarity between our results and the behavior of random Lotka-Volterra models. Servan *et al*.^41^ showed that when growth rates and interaction coefficients are sampled from symmetric distributions with mean zero, the number of coexisting species from a pool of size *n* is also distributed as *B*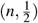. This result holds assuming species are strongly self-limited, such that any biologically feasible equilibrium is stable, as in the CC trade-off model. However, these cases are otherwise very dissimilar. Servan *et al*. considered a scenario where all species are statistically equivalent and interactions are essentially unstructured, while we consider metacommunities with strongly hierarchical interactions, and therefore clear differences between species at opposite ends of the trade-off axis. That the statistics of assembly are identical in these dissimilar cases suggests the intriguing possibility that this is a more general outcome of ecological dynamics where species possess some kind of symmetry, either imposed by assumption, or emerging from ecologically plausible mechanisms, as we demonstrate here.

## Methods

### Model and probabilistic approach

We study a classic metacommunity model, often called the Hastings-Tilman model^15,17^, representing a set of species competing for a large number of habitat patches. This model assumes there is a trade-off between colonization and competitive ability, and that competition is strictly hierarchical; thus, each species *i* is characterized only by its colonization rate (*c*_*i*_) and local extinction rate (*m*_*i*_). The general form of this model was introduced by Tilman^17^:

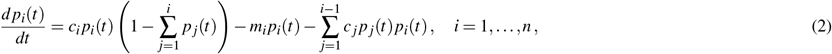

where *p*_*i*_(*t*)∈ [0, 1] tracks the fraction of patches occupied by species *i* at a given time *t* (the occupancy of species *i*), *m*_*i*_ *>* 0 is the local extinction rate of species *i*, and *c*_*i*_ *>* 0 is the colonization rate of species *i*, where species are arranged in increasing order of *c*, such that *c*_1_ *<· · · < c*_*n*_. In other words, *c*_1_ is the colonization rate of the best competitor and *c*_*n*_ is the colonization rate of the weakest.

The first term of Eq. 2 represents the colonization of empty patches by species *i*, the second term represents local extinction (due to frequency-independent factors) in patches where species *i* is currently present, and the last term represents loss of patches due to colonization by superior competitors (i.e. displacement). Each species *i* is always outcompeted (displaced) by species *j < i*, following the competitive hierarchy between species. In this study, we focus on a simplified version of this model where the local extinction rate, *m*, is the same for all species. This assumption can be motivated by regarding *m* as a rate of disturbance that affects all species equally^15^. We also note that this model is a special case of the more general CC trade-off model introduced by Calcagno *et al*.^12^, where they assume that competition is not perfectly asymmetric.

We investigate the metacommunities that emerge through the dynamics of Eq. 2 when the model parameters (*c* values) are sampled at random. More precisely, we take the colonization rates to be a sorted, independent, identically-distributed (*iid*) random sample from a continuous distribution. Sorting, in this context, reflects our assumption of a competition-colonization trade-off. In the language of probability theory, these random colonization rates are *order statistics*^60^.

Treating species’ traits as random samples is a way to uncover model outcomes that are typical and robust, and therefore most likely to be relevant in natural systems. Since this approach was introduced in ecology by the seminal work of May^40^, it has proven to be a productive way to characterize ecological models with many species and parameters, leading to general insights where neither precise empirical determination of parameters nor systematic exploration of parameter space are tractable options^39,61^. Additionally, we go beyond classic applications of this random-interaction approach, which focused on computing properties of a particular equilibrium (e.g. the equilibrium with all *n* species), to study the set of persistent species that are actually assembled through the dynamics of Eq. 2, possibly following the extinction of some species from the pool^41,44–46^.

We note that while drawing traits at random allows us to study outcomes that are independent of the exact values of the model parameters, our conclusions might still depend on the distribution of these parameters, an issue previously discussed by Adler^62^ and Calcagno *et al*.^63^. To maintain generality, we consider an arbitrary continuous distribution with density *f* (*x*), subject only to the assumption that *f* (*c*) = 0 for *c < m*. Species with *c < m* would go extinct even in isolation, so this assumption narrows our focus to a pool of potentially coexisting species – thus, we concentrate on metacommunities where coexistence is determined by biotic interactions, not environmental filtering. For analytical tractability, we consider specific choices of *f* (*x*) for some of our results, although we treat more general cases in SI Section 2 and we explore a range of possible distributions using numerical simulations.

### Equilibria and global stability

Many properties of Eq. 2 can be found by noticing that this model is a special case of the well-known generalized Lotka-Volterra (GLV) model. In particular, we can re-write Eq. 2 in the GLV form

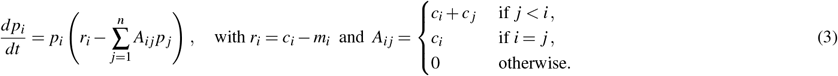

This is a highly structured GLV system where the matrix of interactions, *A*, is lower triangular, corresponding to hierarchical competition, and growth rates, *r* = (*r*_1_, …, *r*_*n*_), are tied to interaction strengths through the colonization rate parameters. Due to our assumption that *c*_*i*_ *> m*_*i*_ for all species, all growth rates are positive.

To study the behaviour of *p*(*t*) = (*p*_1_(*t*), …, *p*_*n*_(*t*)) as *t* → ∞, we characterize the fixed points of the system Eq. 3. Assuming that no two species have identical colonization rates, *A* is an invertible matrix, and a potential fixed point *p*^⋆^ can be explicitly determined by *p*^⋆^ = *A*^−1^*r*. However, this point will only be a fixed point of the dynamics if 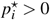 for all *i*. More generally, we are interested in finding subsystems where some species may vanish (i.e. 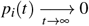 for some *i*) and all those that remain coexist at positive occupancy. For GLV models, there is one potential fixed point corresponding to each unique combination of persisting/extinct species. Moreover, using standard results for the GLV model, it is straightforward to prove that exactly one of these fixed points is *globally stable*, meaning that for every *p*_0_ *>* 0, the solution to Eq. 3 which starts at *p*(0) = *p*_0_ approaches this fixed point as *t* → ∞. The proof of this result, which we present in SI Section 1, relies on the triangularity and non-negativity of *A*.

This unique globally stable equilibrium can be found using a simple algorithm, which also takes advantage of the triangular structure of *A*. Because the dynamics of species *i* only depend on species *j* ≤*i*, the relevant fixed point can be found by setting 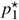 at its equilibrium value, and then computing the equilibrium occupancy for species 2. If this value is positive, it is added to the fixed point; if it is negative, 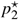 is set to zero. This is repeated for species 3, and so on. This procedure ensures that the resulting equilibrium is both *feasible*, meaning that all species have non-negative occupancy, and *non-invasible*, meaning that extinct species would not be able to re-invade (i.e. they have negative invasion growth rates). We use this algorithm (see SI Section 1) to efficiently find the persisting set of species in our simulations, bypassing the need to numerically integrate the model dynamics. Simulations were conducted in R (version 3.6.3); all code is available at https://github.com/zacharyrmiller/coexistence_random_tradeoff.

### Approximating the persistent set when *n* is large

For a pool of *n* species with colonization rates sampled from some distribution with density *f* (*x*), we aim to calculate the probability that any particular set of species constitutes the unique equilibrium metacommunity, which we denote by ℐ. It is then possible to derive the distribution of key quantities, such as the richness of the final coexisting metacommunity (*S* = |*ℐ* |). ^17^

To calculate these probabilities, we analyze the iterative process described above mathematically. Tilman derived a formula for the *niche shadow* associated with species *i*; that is, a range *c*_*i*_ *< c* _*j*_ *< ℓ*_*i*_ within which any inferior competitor *j* is excluded. Thus, rather than computing 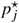 to determine if species *j* is in the coexisting metacommunity (using the algorithm outlined above), one can equivalently compute *ℓ*_*i*_ for each coexisting species and then check whether *c* _*j*_ falls within this niche shadow. This alternative perspective is convenient, because it can be reduced to a recurrence formula. We show (SI Section 1) that the thresholds *ℓ*_*i*_ satisfy

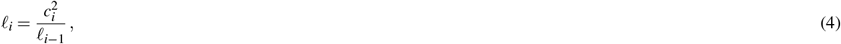

with *ℓ*_0_ = *m*.

Now we introduce the random variable *X*_*i*_ describing the amount by which *c*_*i*_ exceeds its threshold (i.e. *X*_*i*_ = *c*_*i*_− *ℓ*_*i*−1_). If *n* is sufficiently large, *X*_*i*_ will typically be small. We can then use the following approximation for Eq. 4

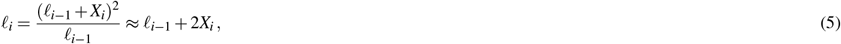

where we neglect the small quadratic term 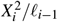. We use this approximation to calculate the probability that a given number of species fall in the interval (*c*_*i*_, *ℓ*_*i*_) ≈ (*c*_*i*_, 2*c*_*i*_ −*ℓ*_*i*−1_).

### Distribution of *S*: elements of the proof

As a first step toward calculating the probability that a given set of species forms the equilibrium metacommunity (and the rest go extinct), we calculate the probability that species 1 excludes the subsequent *k* species in the pool. We define the random variable *K*_1_ to be the number of species excluded by species 1.

Supposing that *c*_1_ ∈ (*x, x* + *dx*), which occurs with probability *f* (*x*) *dx*, the (approximate) niche shadow cast by species 1 extends from *x* to 2*x*−*m*. Conditioning on this value for *c*_1_, the probability that there are exactly *k* species with colonization rates in (*x*, 2*x*−*m*) and *n*−*k* − 1 with colonization rates greater than 2*x*−*m* is given by

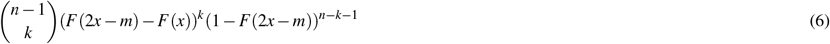

using the assumption that colonization rates are sampled independently. Here *F*(*x*) is the cumulative distribution function associated with *f* (*x*) (i.e. *F*^*′*^(*x*) = *f* (*x*)). The combinatorial factor counts the number of ways to choose *k* excluded species from among the *n* −1 that are not the first.

Multiplying these (independent) probabilities together and integrating over possible values of *x*, we can compute the marginal probability that the first species excludes exactly *k* others:

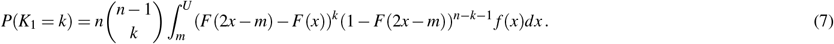

Another combinatorial factor *n* appears because any of the *n* species might be selected as the best competitor (species 1). The integral runs from *m* to *U*, an upper limit that depends on the support of *f* (*x*) and value of *k*. If the distribution of colonization rates is restricted to a finite interval (*m, b*) and *k < n*− 1, then 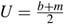, ensuring that the niche shadow does not exceed *b*.

Otherwise *U* = *b* or *U* = ∞, depending on whether the distribution is bounded or not.

In SI Section 2.5, we discuss a large *n* approximation for Eq. 7 when the distribution is arbitrary. Here, for simplicity, we restrict our attention to the Uniform distribution 𝒰 [*m, b*]. Applying the corresponding definitions for *f* and *F*, and using the change of variables 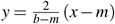, Eq. 7 reduces to

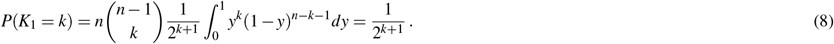

This last equality comes from recognizing the integral as a beta function (see SI Section 2.2 for details). We have assumed here that *k < n*− 1 (and set 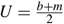 accordingly). In the case *k* = *n*− 1, we show (SI Section 2.2) that the value is 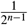 instead.

This result immediately allows us to compute the probability of observing an entire equilibrium configuration. Using the Markov property of order statistics^60^, once we condition on the first *k* + 1 species falling below a certain point (the threshold value *ℓ*_1_ = 2*x* −*m*), and the remaining species above, the distribution of these *n*− *k* −1 remaining colonization rates is independent of the first *k* + 1, and *iid* from the original distribution truncated at *ℓ*_1_. For the Uniform distribution, this truncated distribution is again Uniform. Thus, with very minor modifications, the probability that the second persistent species excludes exactly *k*_2_ species is calculated exactly as above. Continuing this process, and relying on the Markov property to guarantee conditional independence at each step, we can calculate the probability of any particular set of persistent species, specified by the number of excluded species between them (*k*_1_, *k*_2_, …), simply by multiplying. Extend our earlier notation, we define *K*_*i*_ as the number of species excluded by the *i*th persisting species. For example, from a pool of seven species, the coexisting set *{*1, 2, 5*}* would correspond to *K*_1_ = 0, *K*_2_ = 2, and *K*_3_ = 2. The probability of a particular final set of species thus specified is

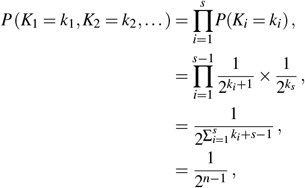

where *s* is the number of persisting species. The final equality follows from the fact that 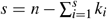 by definition. Remarkably, the final probability does not depend on the particular set of persistent species, or even on *s*. It is equally probable to observe any set of species, so the probability of finding *s* species in the assembled metacommunity is directly proportional to the number of sets of size *s*. More precisely, since the first species always persists, we have

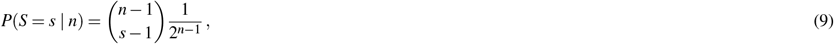

which is the binomial distribution 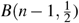 for the number of persisting species in addition to the first (i.e. the random variable *S* − 1 is binomially distributed). From this fact, we can immediately conclude that the average number of persisting species is 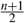, very close to half of the pool.

## Data, script, and code availability

Code to reproduce all simulations and analysis is available at: https://github.com/zacharyrmiller/coexistence_random_tradeoff (DOI: 10.5281/zenodo.7786661).

## Acknowledgements and funding

We thank Vincent Calcagno and Pablo Lechón-Alonso for helpful comments and discussion. J. Timothy Wootton helped inspire this work. Z.R.M. was supported by the National Science Foundation Graduate Research Fellowship Program (DGE-1746045). M.C. and F.M. are supported by the CNRS 80 prime project KARATE.

## Author contributions statement

Z.R.M., M.C., F.M., and S.A. conceived of the research. All authors contributed to the model analysis. Z.R.M. wrote the code for simulations and visualizations. Z.R.M. and M.C. drafted the manuscript. All authors contributed to revising the manuscript.

