## Supplementary Information for "Coexistence of many species under a random competition-colonization trade-off"

July 13, 2023

#### S1 Model properties

The competition-colonization trade-off model (CC trade-off model) can be written as

$$\frac{dp_i(t)}{dt} = c_i p_i(t)\left(1 - \sum_{j=1}^{i} p_j(t)\right) - m_i p_i(t) - \left(\sum_{j=1}^{i-1} c_j p_j(t) p_i(t)\right), \; i = 1, \ldots, n, \tag{1}$$

where $p_i(t) \in [0,1]$ corresponds to the occupancy of species $i$ at a given time $t$ (i.e., the proportion of patches occupied by species $i$ at time $t$), $c_i \in \mathbb{R}_+^\star$ is its colonization rate, and $m_i \in \mathbb{R}_+$ its extinction rate. Species are arranged in increasing order of colonization rate $c$, i.e. $c_1 < \cdots < c_n$. For more details and a derivation of Eq. 1, see Tilman [1].

To motivate our analysis, consider the two-species version of Eq. 1, where species 1 is the best competitor, species 2 the worst competitor, and $m_1 = m_2 = m$. The system's dynamics is governed by two equations:

$$\begin{aligned} \frac{dp_1(t)}{dt} &= c_1 p_1(t)(1 - p_1(t)) - m p_1(t), \\ \frac{dp_2(t)}{dt} &= c_2 p_2(t)(1 - p_1(t) - p_2(t)) - m p_2(t) - c_1 p_1(t) p_2(t). \end{aligned}$$

At equilibrium, the occupancy of species 1 is

$$p_1^\star = 1 - \frac{m}{c_1},$$

which corresponds to the fixed point value of Levins' metapopulation model [2]; note that the equilibrium is feasible whenever $c_1 > m$. For species 2, the occupancy is

$$p_2^\star = 1 - \frac{m}{c_2} - \left(1 + \frac{c_1}{c_2}\right) p_1^\star.$$

This value is lower than the corresponding Levins' model fixed point value, owing to the competitive effect of species 1. Thus, in order to persist, $c_2$ must exceed some value that is larger than $m$. Using the two equilibrium equations, we can deduce the constraints of coexistence between the species

$$c_1 > m \text{ and } c_2 > \frac{c_1^2}{m}.$$

Thus, the coexistence between species depends on the colonization rates meeting certain conditions; however, these conditions are relatively simple. If these rates are considered as random variables, the choice of different distributions of the colonization rates $c$ may result in a different probability of obtaining an admissible (coexisting) pair $(c_1, c_2)$. Here we describe the impact of the distribution $c$ on the coexistence of a pool of $n$ species following Eq. 1.

#### S1.1 Set of admissible solutions

At this point, without concern for the dynamics of Eq. 1, many properties of the model can be obtained in a simple form (see also Tilman [1]). In this section, we first characterize the set of colonization rates $c$ that allows for the persistence of all species, and we show that these conditions are very stringent. Next, we show that the hierarchical nature of the models allow to determine the set of coexisting species in linear time (i.e. performing order $n$ operations; this is in stark contrast with for example the Generalized Lotka-Volterra model, which in general requires computing $2^n$ configurations).

Assume the extinction rates are all equal, i.e. $m_i = m$ for $i = 1, \ldots, n$; then the conditions for species coexistence mainly relate to the choice of the colonization rates $c$ (as seen earlier in the two-species case). Here, the equilibrium occupancy of each species $p^\star = (p_1^\star, ..., p_n^\star)$ can be computed with iterative equations, as well as the condition on the colonization rates to ensure coexistence between species.

First, a rearrangement of Eq. 1 gives

$$\frac{dp_i}{dt} = p_i \left( c_i - m - c_i \sum_{j=1}^{i} p_j - \sum_{j=1}^{i-1} c_j p_j \right) .$$

The equation that gives the non-trivial equilibrium ($p_i^\star > 0$) can be written as follows

$$\frac{dp_i}{dt} = 0 \quad \Leftrightarrow \quad 1 - \frac{m}{c_i} - \sum_{j=1}^{i} p_j - \sum_{j=1}^{i-1} \frac{c_j p_j}{c_i} = 0 \,. \tag{2}$$

One way to solve this equation in the case where $p_i > 0, \ i = 1, \ldots, n$ is to introduce the fraction of patches unoccupied by species $1, \ldots, i$ at equilibrium, defined by

$$h_i^\star = 1 - \sum_{j=1}^{i} p_j \,. \tag{3}$$

Eq. 2 can then be rewritten as

$$-m + c_i h_{i-1}^\star - c_i p_i^\star - \sum_{j=1}^{i-1} c_j p_j^\star = 0 \,. \tag{4}$$

For the iteration $i-1$, we can replace $c_{i-1} h_{i-2}^\star - c_{i-1} p_{i-1}^\star$ by $c_{i-1} h_{i-1}^\star$, which gives

$$-m + c_{i-1} h_{i-1}^\star - \sum_{j=1}^{i-2} c_j p_j^\star = 0. \tag{5}$$

We then remove Eq. 4 from Eq. 5 to obtain a recurrence equation linking $p^\star$ and $h^\star$

$$p_i^\star = \left( 1 - \frac{c_{i-1}}{c_i} \right) h_{i-1}^\star - \frac{c_{i-1}}{c_i} p_{i-1}^\star \,, \tag{6}$$

then we can complete using Eq. 3 rewritten

$$h_i^\star = h_{i-1}^\star - p_i^\star = \frac{c_{i-1}}{c_i}(h_{i-1}^\star + p_{i-1}^\star)\,. \tag{7}$$

Eq. 6 and Eq. 7 can be written as vector equations using the following notations

$$X_i = \begin{pmatrix} p_i^\star \\ h_i^\star \end{pmatrix}, \quad A = \begin{pmatrix} -1 & -1 \\ 1 & 1 \end{pmatrix}, \quad B = \begin{pmatrix} 0 & 1 \\ 0 & 0 \end{pmatrix},$$

Eq. 6 and Eq. 7 are then expressed as

$$X_i = \left(\frac{c_{i-1}}{c_i}A + B\right) X_{i-1}. \tag{8}$$

Suppose we have

$$X_{2i+1} = (\alpha_i A + \beta_i B) X_0, \tag{9}$$

with $p_0 = 0$ and $h_0^\star = 1$. Applying twice in a row the recurrence (8) and the simple computations, we obtain

$$X_{2i+3} = \left(\frac{c_{2i+2}}{c_{2i+3}}A + B\right) X_{2i+2} = \left(\frac{c_{2i+2}}{c_{2i+3}}AB + \frac{c_{2i+1}}{c_{2i+2}}BA\right) X_{2i+1}\,. \tag{10}$$

By injecting Eq. 9 into Eq. 10 and using the relation $ABA = A$, $BAB = B$, we find that

$$X_{2i+3} = \left(\frac{c_{2i+2}}{c_{2i+3}}AB + \frac{c_{2i+1}}{c_{2i+2}}BA\right)(\alpha_i A + \beta_i B)\, X_0 = \left(\frac{c_{2i+2}}{c_{2i+3}}\alpha_i A + \frac{c_{2i+1}}{c_{2i+2}}\beta_i B\right) X_0\,. \tag{11}$$

That is, the following recurrence relations:

$$\alpha_{i+1} = \frac{c_{2i+2}}{c_{2i+3}}\alpha_i\,, \tag{12}$$

$$\beta_{i+1} = \frac{c_{2i+1}}{c_{2i+2}}\beta_i\,. \tag{13}$$

Given $\alpha_0 = m/c_1$, $c_0 = m$ and $\beta_0 = 1$, we derive the general expressions of the coefficients for $i > 0$

$$\alpha_i = \frac{\prod_{j=0}^{i} c_{2j}}{\prod_{j=0}^{i} c_{2j+1}}\,,$$

$$\beta_i = \frac{\prod_{j=0}^{i-1} c_{2j+1}}{\prod_{j=1}^{i} c_{2j}}\,.$$

In the same way, if we suppose

$$X_{2i} = (\gamma_i A + \delta_i B)\, X_1, \quad p_1^\star = 1 - m/c_1, \quad h_1^\star = m/c_1\,.$$

The same method of calculation shows

$$\begin{aligned} X_{2i+2} &= \left(\frac{c_{2i+1}}{c_{2i+2}}A + B\right) X_{2i+1}, \\ &= \left(\frac{c_{2i+1}}{c_{2i+2}}AB + \frac{c_{2i}}{c_{2i+1}}BA\right) X_{2i}, \\ &= \left(\frac{c_{2i+1}}{c_{2i+2}}\gamma_i A + \frac{c_{2i}}{c_{2i+1}}\delta_i B\right) X_1. \end{aligned}$$

This leads to the fact that

$$\gamma_i = \frac{\prod_{j=0}^{i-1} c_{2j+1}}{\prod_{j=1}^{i} c_{2j}}\,, \tag{14}$$

$$\delta_i = \frac{\prod_{j=1}^{i-1} c_{2j}}{\prod_{j=1}^{i-1} c_{2j+1}}\,. \tag{15}$$

It has therefore been shown, with Eq. 12-13 and Eq. 14-15, that the recurrence in Eq. 8 can be decomposed and solved between even and odd indices as follows

$$X_{2i+1} = \left( \frac{\prod_{j=0}^{i} c_{2j}}{\prod_{j=0}^{i} c_{2j+1}} A + \frac{\prod_{j=0}^{i-1} c_{2j+1}}{\prod_{j=1}^{i} c_{2j}} B \right) \cdot \begin{pmatrix} 0 \\ 1 \end{pmatrix}\,, \tag{16}$$

$$X_{2i} = \left( \frac{\prod_{j=0}^{i-1} c_{2j+1}}{\prod_{j=1}^{i} c_{2j}} A + \frac{\prod_{j=1}^{i-1} c_{2j}}{\prod_{j=1}^{i-1} c_{2j+1}} B \right) \cdot \begin{pmatrix} 1 - \frac{m}{c_1} \\ \frac{m}{c_1} \end{pmatrix}\,. \tag{17}$$

In terms of $p_i^\star$, this yields

$$p_{2i+1}^\star = \frac{\prod_{j=0}^{i-1} c_{2j+1}}{\prod_{j=1}^{i} c_{2j}} - \frac{\prod_{j=0}^{i} c_{2j}}{\prod_{j=0}^{i} c_{2j+1}}\,, \tag{18}$$

$$p_{2i}^\star = \frac{m}{c_1} \frac{\prod_{j=1}^{i-1} c_{2j}}{\prod_{j=1}^{i-1} c_{2j+1}} - \frac{\prod_{j=0}^{i-1} c_{2j+1}}{\prod_{j=1}^{i} c_{2j}} = \frac{\prod_{j=0}^{i-1} c_{2j}}{\prod_{j=0}^{i-1} c_{2j+1}} - \frac{\prod_{j=0}^{i-1} c_{2j+1}}{\prod_{j=1}^{i} c_{2j}}\,. \tag{19}$$

It may also be noted that Eq. 16 and Eq. 17 provide expressions for $h_i^\star$ (the unoccupied habitat by species 1 to $i$) at equilibrium. As long as all species persist:

$$h_{2i+1}^\star = \frac{\prod_{j=0}^{i} c_{2j}}{\prod_{j=0}^{i} c_{2j+1}}\,, \tag{20}$$

$$h_{2i}^\star = \frac{\prod_{j=0}^{i-1} c_{2j+1}}{\prod_{j=1}^{i} c_{2j}}\,. \tag{21}$$

By manipulating these two equations, we see that: $h_i^\star = \left(\frac{c_{i-1}}{c_i}\right) h_{i-2}^\star$, which gives a decrease rate of $h_i^\star$ for every two species added.

For species $i$ to persist in the system, its equilibrium given by Eq. 18 or Eq. 19 must be positive. This implies that the conditions for species persistence can be given as

$$p^\star_{2i+1} \text{ persist } \Leftrightarrow c_{2i+1} > m\left(\prod_{j=1}^{i} c_{2j}\right)^2 / \left(\prod_{j=0}^{i-1} c_{2j+1}\right)^2 , \tag{22}$$

$$p^\star_{2i} \text{ persist } \Leftrightarrow c_{2i} > \left(\prod_{j=0}^{i-1} c_{2j+1}\right)^2 / \left[m\left(\prod_{j=1}^{i-1} c_{2j}\right)^2\right] . \tag{23}$$

*Remark* 1. We can reorder the inequalities in Eq. 22-23, so that the conditions of coexistence can be written as ($c_0 = m$)

$$1 < \frac{c_1}{c_0} < \frac{c_2}{c_1} < \frac{c_1}{c_0}\frac{c_3}{c_2} < \frac{c_2}{c_1}\frac{c_4}{c_3} < \frac{c_1}{c_0}\frac{c_3}{c_2}\frac{c_5}{c_4} < ...$$

stopping inequalities with on the right a fraction where the largest index of the numerator is equal to the index of the last persistent species.

*Remark* 2. From Eq. 22-23, it is easy to see that each inequality can be written in a simple recursive form. Let $\ell_i$ be the threshold that $c_i$ must exceed for persistence. Then we have

$$\ell_i = \frac{c_i^2}{\ell_{i-1}} \tag{24}$$

obtained by substituting Eq. 23 into Eq. 22 (or vice versa). We rely on this compact recursive form of the coexistence conditions to derive our main results in Section 2.

Finally the conditions on the colonization rate vector $c$ are formulated in order to have coexistence between the species in the form of a set. The set of admissible solutions represents a series of algebraic conditions and depends on $c$ and $m$

$$\mathcal{C}_m = \left\{ x \in \mathbb{R}^n_+ : x_{2i} > \frac{\left(\prod_{j=1}^{i-1} x_{2j+1}\right)^2}{m\left(\prod_{j=1}^{i-1} x_{2j}\right)^2} \quad ; \quad x_{2i+1} > \frac{m\left(\prod_{j=1}^{i} x_{2j}\right)^2}{\left(\prod_{j=1}^{i-1} x_{2j+1}\right)^2} \right\} . \tag{25}$$

To obtain a large coexisting metacommunity in the CC trade-off model, one could just choose a vector of colonization rates satisfying $c \in \mathcal{C}_m$, corresponding to the case where all species coexist and persist. However, this condition is very stringent. Choosing $c$ this way implies a high degree of "fine-tuning". Thus, we are interested in understanding when a subset of species from a large pool will fulfill the conditions $\mathcal{C}_m$.

Now consider a species pool of $n$ species in which the colonization rates are randomly drawn from a given distribution. Let $\mathcal{S} \subset [n]$ be the subset of indices of the persisting species. The components of the equilibrium vector $p^\star$ can be classified into two classes: the persisting species, $i \in \mathcal{S}$ such that $p^\star_i > 0$ and the vanishing species, $i \in \mathcal{S}^c$ such that $p^\star_i = 0$. Denote by $S = \mid \mathcal{S} \mid$ the random variable representing the number of persisting species. The coexistence problem can be reinterpreted as finding the distribution of the number of persisting species with colonization rates taken from a given distribution i.e. $P(S = k \mid n)$.

From a deterministic point of view, this problem is equivalent to checking $2^n$ possibilities, either the presence or the absence of the species. However, since the model includes a competitive hierarchy, if the first species has the opportunity to invade, it cannot be displaced by the next species. The most competitive species will always have priority. We end up testing only $n$ conditions by following a decision tree (see Figure S1).

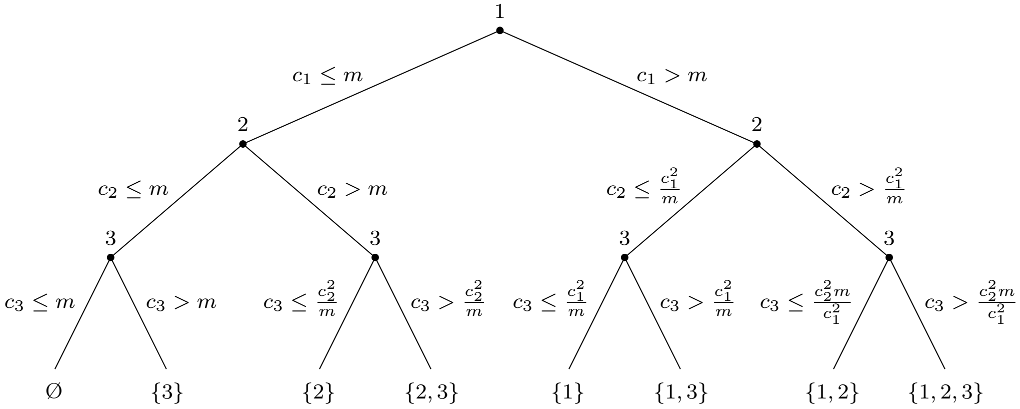


Figure S1: Decision tree for a 3-species system. The path of the binary tree selects the persisting species. At each node, the path on the right corresponds to the persistence of the species, the one on the left to the extinction of the species. At the bottom, the leaves indicate the indexes of the persisting species.

To sum up, the system follows the dynamics given by Eq. 1 starting from a pool of $n$ species. The vector of colonization rates $c$ is sampled from a positive probability distribution and we sort the $c$ in increasing order. The aim is to assess the distribution of the persisting species for a fixed $n$. From an algorithmic standpoint, we traverse the decision tree illustrated in Fig. S1, following the pseudo-algorithm 1 (below) which keeps the persisting species at the equilibrium point.

**Algorithm 1** "All-at-once" scenario

**Require:** $n \geq 0$
  $c \leftarrow$ list
  **for** $l \in [1, n]$ **do** ▷ Creation of a random vector
    randomly choose $c_{new}$ from a probability distribution;
    $c \leftarrow [c, c_{new}]$;
  **end for**
  $c \leftarrow Sorted(c)$; ▷ Ascending sorting algorithm
  **for** $j \in [1, len(c)]$ **do** ▷ Selection by the tree of the persisting species
    $S \leftarrow m$;
    **if** $c[j] \leq S$ **then**
      $del(c[j])$;
    **else**
      $S \leftarrow c[j]^2/S$;
    **end if**
  **end for**

#### S1.2 Equivalence to the Lotka-Volterra model and global stability of the equilibrium

Introduced at the beginning of the 20th century by Lotka [3] and Volterra [4], the generalized Lotka-Volterra (GLV) model is one of the most popular models in ecology. One of its strengths is its versatility: many models can be related to a GLV model including, in particular, the CC trade-off model. Here we recast the CC trade-off model given by Eq. 1 as a GLV model and give a demonstration that this model possesses a globally stable equilibrium point, which is the unique attractor when all species are present in the initial pool, i.e. $p_i(0) > 0\,,\ \forall i \in [n]$. Notice that Tilman [1] and Hastings [5] also address these points in appendix of their articles.

First, we reformulate the CC trade-off model as a GLV model

$$\frac{dp_i}{dt} = c_i p_i \left(1 - \sum_{j=1}^{i} p_j\right) - m_i p_i - \left(\sum_{j=1}^{i-1} c_j p_j p_i\right)\ ,\ i = 1, \ldots, n\,,$$

$$\Leftrightarrow \frac{dp_i}{dt} = p_i \left(r_i - \sum_{j=1}^{n} A_{ij} p_j\right)\ ,\ i = 1, \ldots, n\,, \tag{26}$$

where

$$r_i = c_i - m_i \quad \text{and} \quad A_{ij} = \begin{cases} c_i + c_j & \text{if } j < i\,, \\ c_i & \text{if } i = j\,, \\ 0 & \text{otherwise.} \end{cases}$$

Generally, $r = (r_1, ..., r_n)^\top$ is understood as a vector of growth rates and corresponds here to the dynamics of each species without interactions. If the colonization rate is superior to the extinction rate, the species persists and grows indefinitely, otherwise, the species vanishes. However, this must be interpreted carefully because $p_i \in [0, 1]$, therefore $r$ cannot be clearly understood without $A = (A_{ij})_{n\times n}$. The matrix $A$ corresponds to a matrix of interactions; it is a competitive interaction matrix because $-A_{ij} < 0$, $\forall\, i, j \in [n]$. The impact of species $j < i$ on species $i$ is $-(c_i + c_j)$. This interaction coefficient depends on $c_j$, such that the higher $c_j$ (equivalently, the closer $c_j$ is to $c_i$), the more strongly species $j$ affects the growth of species $i$.

To study the behavior of $p(t)$ as $t \to \infty$, we characterize the equilibrium of Eq. 26. An equilibrium $p^\star$ is defined as a vector satisfying

$$\frac{dp_i^\star}{dt} = 0 \quad \Leftrightarrow \quad p_i^\star(r_i - (Ap^\star)_i) = 0\,, \quad i = 1, \ldots, n\,.$$

If $A$ is non-singular and a feasible fixed point exists, i.e. $p_i^\star > 0$, $\forall i \in [n]$, then the equilibrium $p^\star$ can be explicitly determined by

$$p^\star = A^{-1} r\,.$$

The condition on the vector $c$ to have all species coexisting is $c \in \mathcal{C}_m$, and as we have shown above this is very restrictive. In general, we consider cases in which there is no feasible equilibrium for the full pool of species. Instead, we focus on a fixed point where some species may vanish, i.e $p_i(t) \xrightarrow[t\to+\infty]{} 0$. In the following, we show that Eq. 26 has a unique globally stable equilibria, i.e. all other equilibrium are not stable.

An equilibrium $p^\star$ for Eq. 26 is globally stable if for every $p_0 > 0$, the solution to Eq. 26 which starts at $p(0) = p_0$ satisfies

$$p(t) \xrightarrow[t\to\infty]{} p^\star\,.$$

To prove that Eq. 26 has a unique globally stable equilibrium, we first characterize $A$. Recall that a square matrix $M$ is said to be a $P$-matrix if and only

1. All principal minors of $M$ are positive

$$\det(M_{\mathcal{I}}) > 0\ , \quad \forall \mathcal{I} \subset [n]\ , \quad M_{\mathcal{I}} = (M_{ij})_{ij\in\mathcal{I}}\ .$$

2. All real eigenvalues of $M$ and its principal submatrices are positive.

The matrix $A$ defined in Eq. 26 is a $P$-matrix. The proof relies on two properties of triangular matrices. Given $A$ is a lower triangular matrix, the eigenvalues are the diagonal entries of the matrix and $c$ is positive. Second, each principal submatrix of $A$ is a lower triangular matrix and its eigenvalues correspond to a subset of $c > 0$. Using the second definition of a $P$-matrix ends the proof.

In graph theory, the matrix $A$ represents a directed acyclic graph. A directed acyclic graph is a directed graph ($A$ is a non-empty directed path in which only the first and last vertices are equal) where there is no cycle in the graph. Consequently, $-A$ is composed of cycles of length one.

A Theorem of Takeuchi *et al.* [6] states that if $-A$ has only cycles of length one, then the system of Eq. 26 and every reduced system of Eq. 26 have a nonnegative and globally stable equilibrium point for each $r \in \mathbb{R}^n$ if and only if $A$ is a $P$-matrix.

To conclude, the system of Eq. 26, equivalent to Eq. 1, has a unique globally stable equilibrium point $p^\star$ independently of the parameter values, i.e. for any positive initial condition $p_0$, colonization rate $c$ and extinction rate $m$. Algorithm 1 yields this unique globally stable equilibrium point for any choice of $c$ and $m$, since this procedure finds a nonnegative equilibrium point that is non-invasible by any of the species with $p_i^\star = 0$.

#### S1.3 Choice and impact of the extinction rate

**Choice of the extinction rate** Here, we focus on a specific CC trade-off model form of Eq. 1 similar to the model studied by Hastings [5]. The extinction rate is equal for each species, i.e. $m_i = m$ for every $i = 1, \ldots, n$. Choosing equal extinction rates reduces the number of parameters and greatly simplifies the complexity of the model. Understanding the impact of the colonization rates $c$ in the CC trade-off model is sufficient to capture many open problems and represent the main interest of this paper. However, studying the model with different extinction rates is also an intriguing perspective (see, e.g., May & Nowak [7]).

**Impact of the extinction rate** The impact of the extinction rate has been studied and introduced by Hastings [5] in the CC trade-off model. The number of species in the system differs according to the extinction rate, a relationship known as the diversity disturbance relationship. In this model, the number of persistent species as a function of the extinction rate has an optimal intermediate value. For any distribution of colonization rates, the number of persisting species reaches a peak when the left support of the probability distribution of the colonization rates is at $m$. Let $c_{\min}$, the left edge of the support of the distribution. In the following, we fix the value $c_{\min} = 1$. At $m = c_{\min}$, we will show in the following sections that the richness converges to the binomial distribution – about half of the species persist independently of the chosen distribution. On the other hand, when $m \neq c_{\min}$, there is a loss of diversity, explained by

- if $m \gg c_{\min}$, a large part of the density of the distribution is truncated with the first condition $c_i > m$, $i = 1, \ldots, n$. Species that have a lower colonization rate than the extinction rate cannot persist.
- if $m \ll c_{\min}$: a large part of the density of the distribution is truncated with the second condition; for example, if we suppose that $c_1$ is very close to $c_{min} = 1$, then $c_2 > \frac{c_1^2}{m} \approx \frac{1}{m}$. Species with colonization rates less than the inverse of the extinction rate cannot persist.

From an ecological point of view, an increased extinction rate of all species is generally associated with a decrease in species richness. However, in this model, a decrease in extinction rate can imply a decrease in species richness. This recalls the well-known intermediate disturbance relationship (see [8]). In the CC trade-off model, a decrease in extinction rate implies a supremacy of the most competitive species at the expense of the others, hence the decrease in richness.

Without loss of generality and in a framework where the interest is on $c$, we could have chosen an extinction rate equal to $m = 1$ for all species. To see this, begin with Eq. 1 where we divide the equation by $m$

$$\frac{dp_i}{dt} = p_i c_i \left(1 - \sum_{j=1}^{i} p_j\right) - m p_i - \left(\sum_{j=1}^{i-1} c_j p_j p_i\right) , \tag{27}$$

$$\Leftrightarrow \quad \frac{dp_i}{dt}\frac{1}{m} = p_i \frac{c_i}{m} \left(1 - \sum_{j=1}^{i} p_j\right) - p_i - \left(\sum_{j=1}^{i-1} \frac{c_j}{m} p_j p_i\right) .$$

One can define a new vector $c'$ where $c'_i = \frac{c_i}{m}$ , $i = 1, \ldots, n$. Considering $m$ fixed, the analyses can proceed equivalently on $c$ or $c'$:

$$\frac{dp_i}{dt}\frac{1}{m} = p_i c'_i \left(1 - \sum_{j=1}^{i} p_j\right) - p_i - \left(\sum_{j=1}^{i-1} c'_j p_j p_i\right) . \tag{28}$$

The equilibrium of Eq. 28 is similar to Eq. 27 with $m = 1$. The factor $1/m$ has only an impact on the speed of the convergence. Denote by $\mathcal{C}'_1$ the series of algebraic conditions associated with the vector $c'$, then

$$c' \in \mathcal{C}'_1 \quad \Leftrightarrow \quad c \in \mathcal{C}_m .$$

However, we will maintain the term $m$ (when possible) for clarity throughout.

### S2 Distribution of richness in the assembled metacommunity

In this section, we examine the distribution of the number of persisting species (i.e. the number of nonzero components in the unique globally stable equilibrium). Calculating this distribution exactly is difficult, even for small $n$. Instead, we focus on characterizing the limiting distribution for large $n$, where we can employ approximations that simplify the mathematical problem. After introducing our key approximation, we derive the distribution of richness for the special cases where the distribution of colonization rates is either Uniform or Exponential. In each case, we adopt the strategy outlined in the Main Text: we calculate the probability of observing any particular set of persisting species by sequentially calculating the probability that the best competitor (the species with the smallest colonization rate in the pool) excludes $k_1$ species, that the next persisting species excludes $k_2$ species, and so on. We also provide a partial characterization for the Triangular distribution, which sheds light on deviations from the binomial theoretical prediction that occur for finite $n$. Finally, using some additional approximations, we show how the binomial distribution arises as the limiting distribution of richness for a wide class of colonization rate distributions.

#### S2.1 Linear niche shadow approximation

To approximate the niche shadow cast by each species, we first express the threshold $\ell_i$ as

$$\ell_i = \frac{c_i^2}{\ell_{i-1}} = \frac{(\ell_{i-1} + X_i)^2}{\ell_{i-1}} = \frac{\ell_{i-1}^2 + 2\ell_{i-1}X_i + X_i^2}{\ell_{i-1}} = \ell_{i-1} + 2X_i + \frac{X_i^2}{\ell_{i-1}} ,$$

where $X_i = c_i - \ell_{i-1}$ is the amount by which $c_i$ exceeds its lower bound ($\ell_{i-1}$). Our approximation will be motivated by the fact that $\ell_{i-1}$ is typically much larger than $X_i$.

To see this, we first note that $\ell_{i-1} \geq \ell_0 = m$. Now we consider $X_i$. As a first approximation, for a distribution of colonization rates with finite support on $(m, b)$, this random variable will typically be on the order $(b - m)/n$. To obtain a better (and more general) approximation, we can study the distribution of $X_i$ when $n$ is large. The conditional (marginal) distribution of the remaining colonization rates, given that the first $k$ (including both the $i-1$ persisting species and the excluded species) are less than $\ell_{i-1}$ and the remaining $n-k$ are greater is

$$F'(c) = \frac{F(c) - F(\ell_{i-1})}{1 - F(\ell_{i-1})} \,.$$

The probability that $X_i < x$ is given by the probability that the minimum among these $n-k$ rates is less than $F(\ell_{i-1} + x)$, which is easily computed as

$$\begin{aligned} P(X_i < x) &= 1 - (1 - F'(\ell_{i-1} + x))^{n-k} \\ &= 1 - \left(1 - \frac{F(\ell_{i-1} + x) - F(\ell_{i-1})}{1 - F(\ell_{i-1})}\right)^{n-k} \,. \end{aligned} \tag{29}$$

Next we note that, for large $n$, $F(\ell_{i-1}) \approx \frac{k}{n}$. This follows from the fact that the number of *iid* rates less than $\ell_{i-1}$ is distributed as $\mathrm{Binom}(n, F(\ell_{i-1}))$, and so $k$ is almost always near $nF(\ell_{i-1})$ for large $n$. We can then approximate the probability in Eq. 29 by

$$P(X_i < x) \approx 1 - \left(1 - \frac{n(F(\ell_{i-1} + x) - F(\ell_{i-1}))}{n-k}\right)^{n-k} \approx 1 - \exp\left(-n(F(\ell_{i-1} + x) - F(\ell_{i-1})\right) \,,$$

where the last approximation holds assuming $n - k$ is also sufficiently large. Finally, we focus our attention on this probability for small $x$. Taylor expanding $F(\ell_{i-1} + x)$ around $\ell_{i-1}$, we obtain

$$P(X_i < x) \approx 1 - \exp(-nf(\ell_{i-1})x) \,. \tag{30}$$

This is the CDF for an Exponential random variable with rate $\lambda = nf(\ell_{i-1})$. We note that as $n \to \infty$, the probability that $X_i$ is very close to zero approaches one (consider, for example, $P(X_i < n^{-1/2})$), implying that the distribution of $X_i$ can be well understood by looking at the small $x$ behavior. Because $X_i$ is well approximated by an Exponential random variable, we conclude that $X_i$ is typically on the order of $1/\lambda = 1/(nf(\ell_{i-1}))$. Thus, provided $n - k$ is large and $f(\ell_{i-1})$ is not too small, $X_i$ is on the order of $1/n$. These assumptions are reasonable outside of the tails of the colonization rate distribution.

Returning to $\ell_i$, we see that $X_i$ is small compared to $\ell_{i-1}$ whenever $nm \gg 1$. Hence, we neglect the term $X_i^2/\ell_{i-1}$, obtaining

$$\ell_i \approx \ell_{i-1} + 2X_i \,, \tag{31}$$

which is Eq. 5 in the Main Text. This is equivalent to $\ell_i \approx 2c_i - \ell_{i-1}$.

We verify the accuracy of this approximation using numerical simulations, as shown in Figs. S2, S3, and S4. Fig. S2 shows that Eq. 30 provides an excellent match to simulations. Figs. S3 and S4 show the resulting approximations for $\ell_i$, obtained by iterating Eq. 31. As we would expect based on the above considerations, this approximation becomes very good for large $n$. We observe that the exact and approximate values of $\ell_i$ diverge substantially only for the largest values of $i$; i.e., in the upper tail of the distribution, as predicted. Notably, even when there are quantitative mismatches between the exact and approximate values, our approximation scheme predicts the number of persisting species with very high accuracy. Thus, we study the recursive process described by Eq. 31 in the following sections to derive the approximate distribution of richness in the assembled metacommunity, for different colonization rate distributions.

It is worth noting that these arguments already suggest that approximately half of the species will persist in the assembled metacommunity. Eq. 31 implies that the niche shadow cast by the

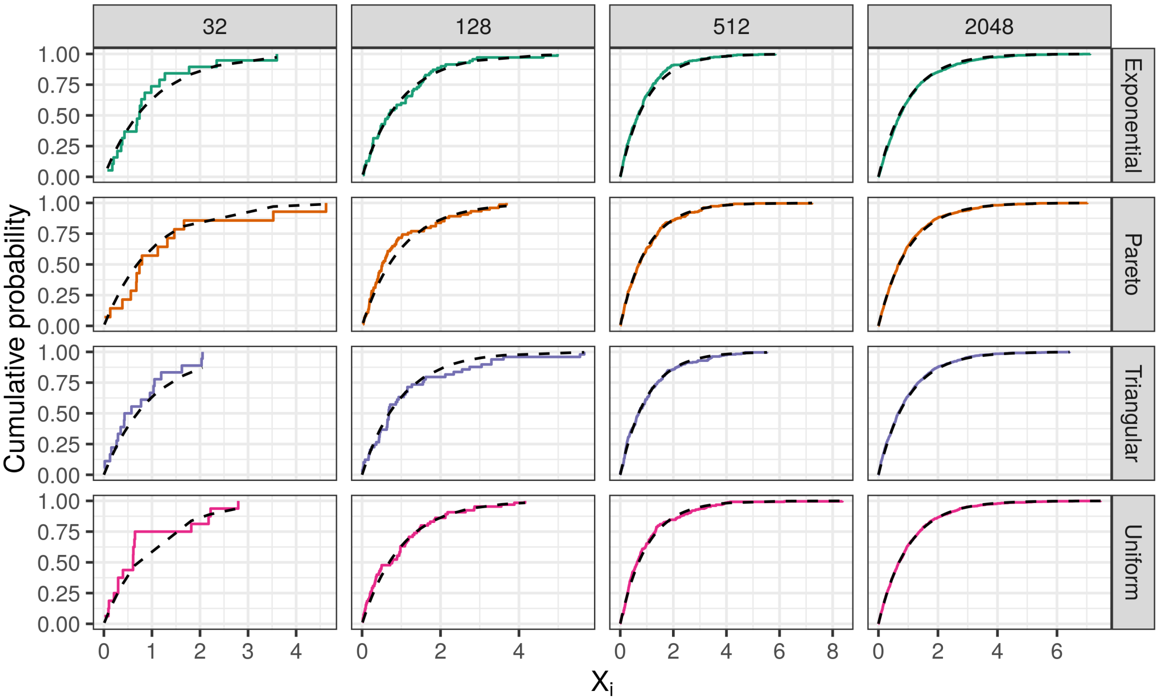


Figure S2: Distribution of $X_i$ is approximately Exponential. Here we plot the empirical cumulative distribution of the (rescaled) quantities $X_i = c_i - \ell_{i-1}$ for all $i$ in simulations with different pool sizes (columns) and colonization rate distributions (rows). Following the approximation theory developed in the text, we rescale each observed $X_i$ by $nf(\ell_{i-1})$; the distribution of rescaled values should collapse onto the Exponential distribution with rate 1 (black dashed line). This approximation is very good, especially for large $n$. Each panel shows results from a single assembled metacommunity with randomly sampled colonization rates.

$i$th persisting species should be approximately equal in length to $X_i$. Thus, the length of this niche shadow is approximately an Exponential random variable with rate $nf(\ell_{i-1})$. By the same logic explained above, the gap between $c_i$ and the next colonization rate in the pool should be distributed approximately as $\text{Exp}(nf(c_i))$. Because $c_i$ and $\ell_{i-1}$ are very close for large $n$, these two random variables – the length of the niche shadow and the gap between $c_i$ and the next colonization rate in the pool – have nearly identical distributions. Thus, by symmetry, the probability that the next species in the pool will be excluded is about one half.

#### S2.2 Uniform distribution

Using Eq. 31, we first consider the simple case where colonization rates are Uniformly distributed between $m$ and some upper limit $b$. As discussed in the Main Text, we can express the probability that the best competitor (i.e. the species with colonization rate $c_1$) excludes the subsequent $k_1$ species as

$$P(K_1 = k_1) = n\binom{n-1}{k_1} \int_m^U (F(2x-m) - F(x))^{k_1} (1 - F(2x-m))^{n-k_1-1} f(x)dx\,, \qquad (32)$$

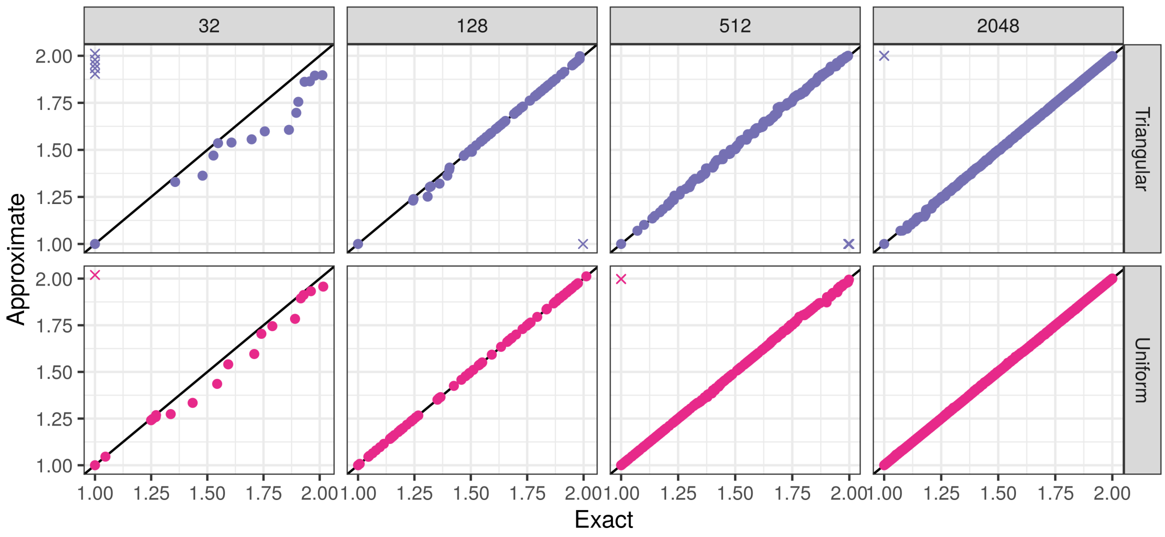


Figure S3: Actual (exact) threshold values $\ell_i$ vs approximate values obtained by iterating Eq. 31 for the Uniform and Triangular distributions. The 1:1 line is shown in black. Where the two predict different numbers of persisting species, the observations (predictions) with no corresponding values are shown along the x- (y-) axis with an "X". For large $n$ we observe very good agreement, and the two processes predict very similar numbers of persisting species.

for any distribution of colonization rates with CDF $F(x)$, PDF $f(x)$, and support on some interval beginning at $m$. Note that the random variable $K_1$ may be zero. This formula uses the approximation that the niche shadow cast by the first species extends from $x$ to $2x - m$.

Specializing to the case of the Uniform distribution on $(m, b)$, we have $F(x) = \frac{x-m}{b-m}$ and $f(x) = \frac{1}{b-m}$ for $x \in (m, b)$. We also consider the upper limit of integration, $U$. For $k_1 < n - 1$, at least one species persists beyond the niche shadow of species 1. This implies that the niche shadow of species 1 falls short of $b$. To accommodate this restriction, we must choose $U = \frac{b+m}{2}$, so that $l_1 \approx b$ at $x = U$. In the case where $k_1 = n - 1$, then the niche shadow may extend beyond $b$. This case must be handled separately, as we discuss below.

Using these definitions, Eq. 32 becomes

$$\begin{aligned}
P(K_1 = k_1) &= n\binom{n-1}{k_1}\int_m^{\frac{b+m}{2}} \left(\frac{2(x-m)}{b-m} - \frac{x-m}{b-m}\right)^{k_1}\left(1 - \frac{2(x-m)}{b-m}\right)^{n-k_1-1}\frac{1}{b-m}dx\,,\\
&= n\binom{n-1}{k_1}\int_m^{\frac{b+m}{2}} \left(\frac{x-m}{b-m}\right)^{k_1}\left(1 - \frac{2(x-m)}{b-m}\right)^{n-k_1-1}\frac{1}{b-m}dx\,.
\end{aligned}$$

To simplify this expression, we introduce the change of variables $y = \frac{2}{b-m}(x - m)$ and obtain

$$\begin{aligned}
P(K_1 = k_1) &= n\binom{n-1}{k_1}\int_0^1 \left(\frac{y}{2}\right)^{k_1}(1-y)^{n-k_1-1}\frac{1}{2}dy\,,\\
&= n\binom{n-1}{k_1}\frac{1}{2^{k_1+1}}\int_0^1 y^{k_1}(1-y)^{n-k_1-1}dy\,.
\end{aligned}$$

We can now recognize this integral as the beta function $B(k_1 + 1, n - k_1)$ [9]. For positive integers $z_1$ and $z_2$, we have the relationship

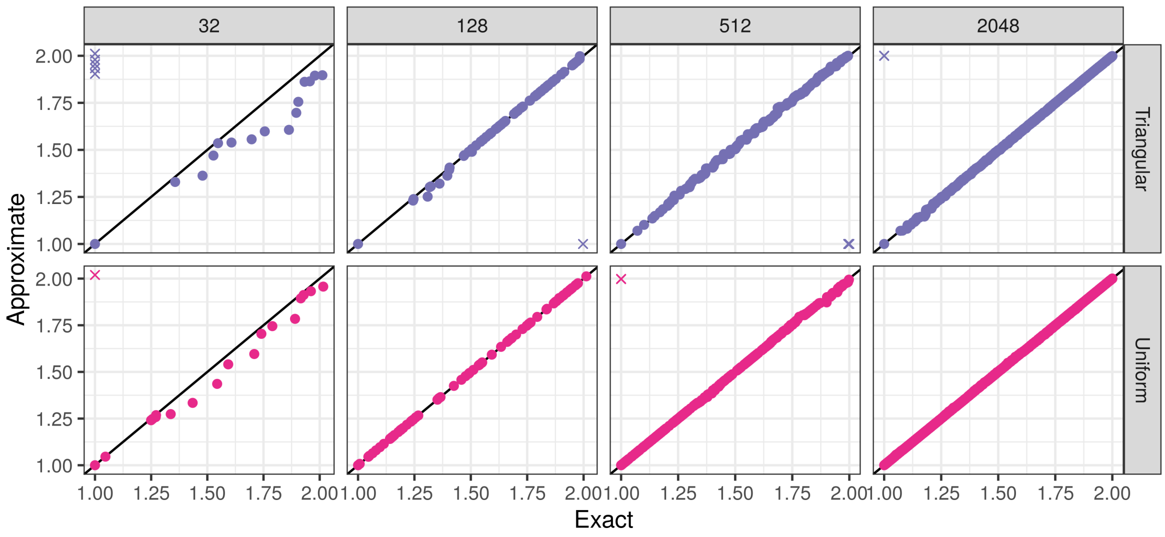


Figure S4: As Fig. S3, but for the Exponential and Pareto distributions. For these distributions, which have unbounded tails, we observe more substantial deviations between the exact and approximate values, but these are largely confined to the tails. Note that for better visualization, we show results on log-log scales.

$$B(z_1, z_2) = \frac{(z_1 - 1)!(z_2 - 1)!}{(z_1 + z_2 - 1)!}\,, \tag{33}$$

and consequently

$$B(k_1 + 1, n - k_1) = \frac{k_1!(n - k_1 - 1)!}{n!} = \frac{1}{n\binom{n-1}{k_1}}\,.$$

Thus, we obtain $P(K_1 = k_1) = \frac{1}{2^{k_1+1}}$ following the cancellation of the combinatorial factors.

Before proceeding, we consider the special case $k_1 = n - 1$. Here we must distinguish two possibilities: (i) the value of $c_1$ is below $U = \frac{b+m}{2}$, and the niche shadow of species 1 falls within the range of colonization rates, or (ii) the value of $c_1$ is between $U$ and $b$, which occurs if all colonization rates in the pool are greater than $U$. In the former case, the probability that $n - 1$ species fall within the niche shadow, and zero outside, is calculated as before. In the latter case, exclusion of the other $n - 1$ species occurs with certainty. Adding together these two mutually exclusive possibilities, we have

$$P(K_1 = n - 1) = n \int_m^{\frac{b+m}{2}} \left(\frac{x - m}{b - m}\right)^{n-1} \frac{1}{b - m} dx + \left(1 - \frac{U - m}{b - m}\right)^n = \frac{1}{2^n} + \frac{1}{2^n} = \frac{1}{2^{n-1}}$$

using the arguments above to evaluate the integral. Now that we have calculated $P(K_1 = k_1)$ for $k_1 = 0, \ldots, n - 1$, it is easy to verify that these probabilities define a valid probability mass function:

$$\sum_{i=0}^{n-1} P(K_1 = i) = \sum_{i=0}^{n-2} \frac{1}{2^{i+1}} + \frac{1}{2^{n-1}} = \sum_{i=1}^{n} \frac{1}{2^i} + \frac{1}{2^n} = 1 - \frac{1}{2^n} + \frac{1}{2^n} = 1$$

using the formula for partial sums of a geometric series.

These calculations are sufficient to derive the probability of observing any set of persisting species. If we condition on the fact that the first $K_1 + 1$ colonization rates fall below $2x - m$, and the rest above, then the Markov property of order statistics [10] implies that these $n - K_1 - 1$ remaining rates are Uniformly distributed on $(2x - m, b)$, independently of the first $K_1 + 1$ rates. Letting $m' = 2x - m$ and $n' = n - K_1 - 1$, it is easy to see that $P(K_2 = k_2)$ – the probability that the second persisting species excludes exactly $k_2$ other species – can be calculated exactly as before. We can iterate this procedure to calculate the probability that $K_3 = k_3$, $K_4 = k_4$, and so on. Any set of persisting species can be specified by the associated sequence of excluded species between them. For $\mathcal{S} = \{1, \sigma_2, \cdots, \sigma_s\}$, the set of indices of the species that constitute the equilibrium metacommunity, we have $k_1 = \sigma_2 - 2, k_2 = \sigma_3 - \sigma_2 - 1, \ldots, k_s = n - \sigma_s$. We note that this implies $\sum_{i=1}^{s} k_i = n - s$, in agreement with the fact that the total number of excluded species must be equal to the total number of species in the pool ($n$) minus the number of persisting species ($s$).

The probability of observing any particular $\mathcal{S}$ can be obtained by straightforward multiplication:

$$
\begin{aligned}
P\left(K_1 = k_1, K_2 = k_2, \ldots\right) &= \prod_{i=1}^{s} P(K_i = k_i)\,, \\
&= \prod_{i=1}^{s-1} \frac{1}{2^{k_i+1}} \times \frac{1}{2^{k_s}}\,, \\
&= \frac{1}{2^{\sum_{i=1}^{s} k_i + s - 1}}\,, \\
&= \frac{1}{2^{n-1}}\,.
\end{aligned}
$$

Note that the $s$th (final) species in the coexisting set necessarily excludes all remaining species; thus we have $P(K_s = k_s) = \frac{1}{2^{k_s}}$, which includes an extra factor of 2, as described above for the case $k_1 = n-1$.

The fact that this probability is the same for all choices of $\mathcal{S}$ makes it easy to calculate the probability of observing a specific number of species in the final metacommunity. There are $\binom{n-1}{s-1}$ distinct communities of size $s$ given a pool of $n$ species (and accounting for the fact that the best competitor always persists). This gives us the distribution

$$
P(S = s \mid n) = \binom{n-1}{s-1} \frac{1}{2^{n-1}}\,. \tag{34}
$$

Thus, the random variable $S - 1$, the number of persisting species in addition to the first, is distributed as $\mathrm{Binom}(n-1, 1/2)$.

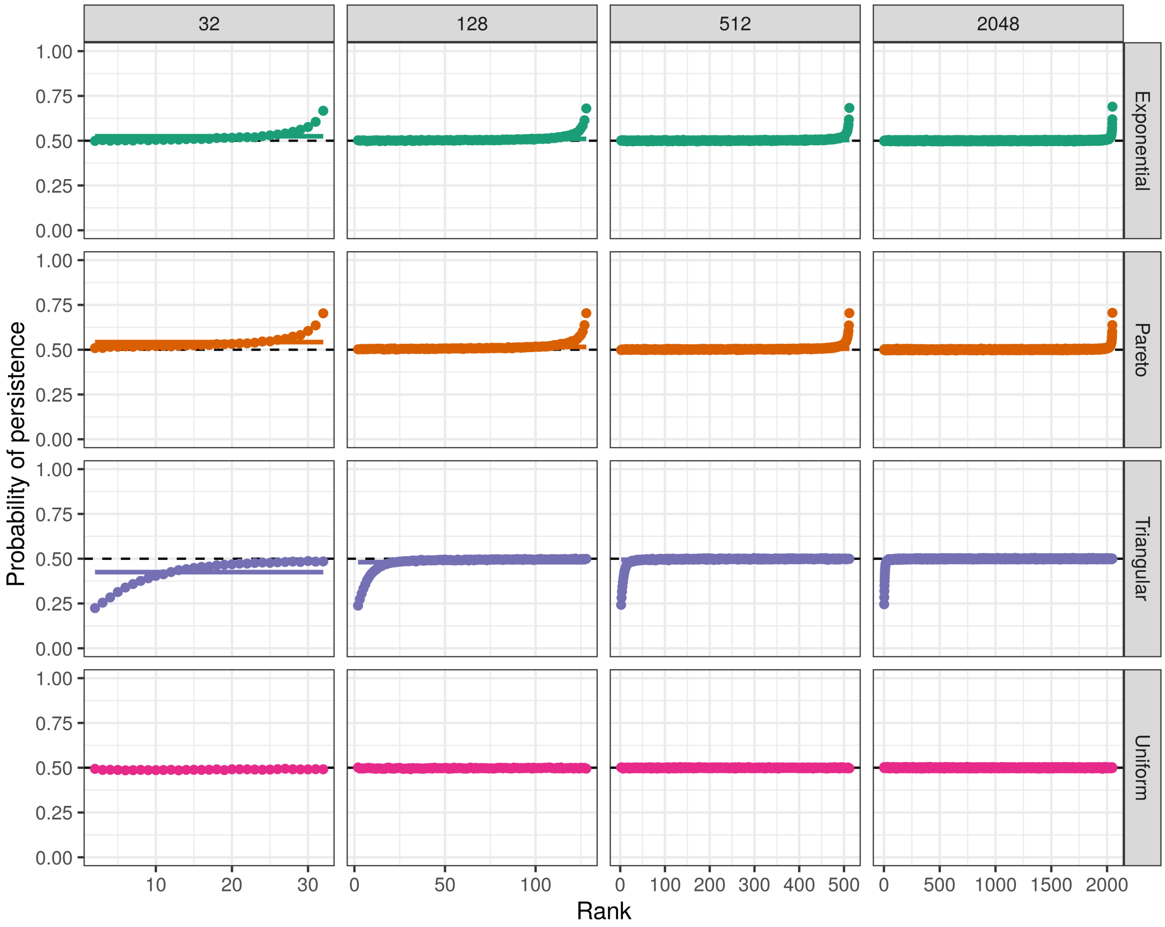


Figure S5: Probability of persisting in the assembled metacommunity by competitive rank in the pool (i.e. species 1 is the best competitor). For each combination of pool size (columns) and colonization rate distribution (rows), the plotted probabilities summarize $10^5$ outcomes from random pools. The marginal probability of persistence, across all ranks, is shown with a solid horizontal line. The theoretical prediction of $\frac{1}{2}$ for all species is shown with a dashed line. Notice predictable deviations for the Exponential and Pareto distributions in the upper tails (best colonizers) and Triangular distribution in the lower tail (best competitors). The fraction of species exhibiting these deviations shrinks as $n$ grows.

#### S2.3 Exponential distribution

We can also evaluate Eq. 32 for the case of an Exponential distribution with rate $\lambda$, shifted so that the distribution has support on $[m, \infty)$. More precisely, in this case we have

$$F(x) = 1 - e^{-\lambda(x-m)} \quad \text{and} \quad f(x) = \lambda e^{-\lambda(x-m)} .$$

Eq. 32 becomes

$$P(K_1 = k_1) = \\ n\binom{n-1}{k_1}\int_m^\infty \left((1-e^{-\lambda(2x-2m)}) - (1-e^{-\lambda(x-m)})\right)^{k_1}\left(1-(1-e^{-\lambda(2x-2m)})\right)^{n-k_1-1}\lambda e^{-\lambda(x-m)}dx\,,$$

or, using the change of variables $y = e^{-\lambda(x-m)}$ (with $dy = -\lambda e^{-\lambda(x-m)}dx$) and simplifying:

$$\begin{aligned} P(K_1 = k_1) &= n\binom{n-1}{k_1}\int_0^1 \left(y-y^2\right)^{k_1}\left(y^2\right)^{n-k_1-1}dy\,, \\ &= n\binom{n-1}{k_1}\int_0^1 (1-y)^{k_1}(y)^{2n-k_1-2}dy\,. \end{aligned} \tag{35}$$

We recognize the integral as a beta function, $B(2n-k_1-1, k_1+1)$ [9]. Using the identity Eq. 33, we find

$$P(K_1 = k_1) = n\binom{n-1}{k_1}\frac{(2n-2-k_1)!\,k_1!}{(2n-1)!} = \frac{n!\,(2n-k_1-2)!}{(n-k_1-1)!\,(2n-1)!} = \prod_{i=0}^{k_1}\frac{n-i}{2(n-i)+(i-1)}\,. \tag{36}$$

In the limit $n \to \infty$, each factor in this product approaches the value $\frac{1}{2}$. Consequently,

$$P(K_1 = k_1) \approx \frac{1}{2^{k_1+1}}\,, \tag{37}$$

exactly as for the Uniform case. As in the Uniform case, we can condition on the first $K_1+1$ rates falling below $2x-m$ and calculate $P(K_2 = k_2)$, $P(K_3 = k_3)$, and so on. Here, we rely on the memoryless property of the Exponential distribution; once we have conditioned on the remaining $n-K_1-1$ rates being greater than $2x-m$, these rates are *iid* samples from an Exponential distribution with the same rate, $\lambda$, shifted so that the conditional CDF is $F'(z) = 1-e^{-\lambda(z-(2x-m))}$. Thus, we can repeat the calculation above for $K_2, K_3, \ldots$, and by the same arguments used for the Uniform distribution, we conclude that all sets of persisting species are about equally likely, and the distribution of $S-1$ becomes approximately $\mathrm{Binom}(n-1, 1/2)$ for large $n$.

For finite values of $n$, Eq. 36 can explain deviations that we observe from the limiting binomial prediction. We observe that the first factor in $P(K_1 = k_1)$, corresponding to $i=0$, is $\frac{n}{2n-1}$, which is greater than $\frac{1}{2}$ for all $n$. The second factor, corresponding to $i=1$, is exactly $\frac{1}{2}$. For $i>1$, each factor is strictly less than $\frac{1}{2}$. This implies that for finite $n$, Eq. 37 underestimates $P(K_1 = k_1)$ for small values of $k_1$ and overestimates it for large values. In particular, one can easily show that $P(K_1 = 0)$ and $P(K_1 = 1)$ are always underestimated, while other probabilities are strictly overestimated. Even when the species pool is large, these discrepancies become important in the tail of the distribution, where the number of species remaining at each step of the calculation becomes small. Among the species with the highest colonization rates, each persisting species is more likely to exclude 0 or 1 other species, leading to "excess" coexistence. For example, in Fig. S5, we plot the probability that each species in the pool is found in the set of persisting species, as a function of its rank colonization (equivalently, competitive) ability. The species with the highest colonization rates are somewhat more likely to be found in the coexisting metacommunity. However, we also observe that this effect is confined to a small number of species in large pools. Thus, the binomial distribution provides a very good approximation for overall richness as $n$ becomes large.

While Eq. 36 holds only for the Exponential distribution, similar deviations appear in the tail of the colonization rate distribution for other right-skewed distributions, such as the Pareto distribution (Fig. S5).

#### S2.4 Triangular distribution

In the preceding sections, we have considered distributions that have high density at $m$, such that there is no left "tail". When $f(m+\epsilon)$ is small for small $\epsilon$, we might expect that the first few species have wider niche shadows, and consequently exclude more competitors. We can see this phenomenon quantitatively by computing $P(K_1 = k_1)$ for the Triangular distribution, which has zero density at $m$ (to simplify the following calculations, we assume support on $(m, m+1)$; an arbitrary upper bound $b$ can easily be treated by rescaling). We have

$$F(x) = (x-m)^2 \quad \text{and} \quad f(x) = 2(x-m)\,.$$

Eq. 32 becomes

$$\begin{aligned} P(K_1 = k_1) &= n\binom{n-1}{k_1}\int_m^U \left((2x-2m)^2-(x-m)^2\right)^{k_1}\left(1-(2x-2m)^2\right)^{n-k_1-1} 2(x-m)dx\,, \\ &= n\binom{n-1}{k_1}\int_m^U \left(4(x-m)^2-(x-m)^2\right)^{k_1}\left(1-4(x-m)^2\right)^{n-k_1-1} 2(x-m)dx\,. \end{aligned}$$

Here we have the upper bound $U = \frac{2m+1}{2}$ for $k_1 < n-1$, as for the Uniform distribution. Now using $y = 4(x-m)^2$ we have

$$P(K_1 = k_1) = n\binom{n-1}{k_1}\int_0^1 \left(\frac{3}{4}y\right)^{k_1}(1-y)^{n-k_1-1}\frac{1}{4}dy\,.$$

Recognizing once again the beta function $B(k_1+1, n-k_1)$, we find

$$P(K_1 = k_1) = \frac{3^{k_1}}{4^{k_1+1}}\,. \tag{38}$$

Eq. 38 suggests that the second species in the pool should persist with probability $\frac{1}{4}$, rather than $\frac{1}{2}$. This is in good agreement with the simulation results shown in Fig. S5. Furthermore, $P(K_1 = k_1)$ is less than the prediction $\frac{1}{2^{k_1+1}}$ for $k_1 = 0$ and $k_1 = 1$, and greater for all other values of $k_1$. This implies that the first species is likely to exclude more competitors than in the cases we have examined above.

Unlike the Uniform and Exponential cases, we cannot simply iterate this calculation to obtain the probability of observing a particular sequence of persisting species. This is because truncating the Triangular distribution at some value $2x-m$ does not produce another Triangular distribution. However, numerical simulations show that the probability of persistence of the $i$th best competitor quickly approaches $\frac{1}{2}$ in large pools, indicating that the distribution of the number of species excluded by each persisting species quickly converges to Eq. 37 once the distribution has been truncated several times. Intuitively, the truncated distribution becomes "flatter" at each step, becoming more and more similar to a Uniform distribution. Because the excess exclusion that we find associated with the first species is confined only to the left tail of the pool, the overall reduction in coexistence associated with the Triangular distribution becomes negligible for large $n$.

#### S2.5 Heuristic approach for arbitrary distributions

Evaluating Eq. 32 is not possible for arbitrary choices of $F(x)$. However, our theoretical and simulation results both suggest that this probability distribution becomes well approximated by Eq. 36 for a wide range of colonization rate distributions when $n$ is large. To obtain insight into this apparently somewhat universal behavior, here we consider a further approximation for $P(K_1 = k_1)$ for arbitrary $F(x)$. We Taylor expand this distribution function around $m$ and use $F(x) \approx F(m) + f(m)(x-m) = f(m)(x-m)$.

This approximation is motivated by our earlier arguments that the gap between $\ell_{i-1}$ (in this case $\ell_{i-1} = m$) and the next colonization rate in the pool is typically very small. Thus, the integral in Eq. 32 will be dominated by the contributions close to $x = m$.

Using this simple linear approximation, Eq. 32 becomes

$$\begin{aligned} P(K_1 = k_1) &\approx n\binom{n-1}{k_1} \int_m^U \left(f(m)(2x-2m) - f(m)(x-m)\right)^{k_1} \left(1 - f(m)(2x-2m)\right)^{n-k_1-1} f(m)dx\,, \\ &= n\binom{n-1}{k_1} \int_m^U \left(f(m)(x-m)\right)^{k_1} \left(1 - 2f(m)(x-m)\right)^{n-k_1-1} f(m)dx\,. \end{aligned}$$

Here we choose $U = m + \frac{1}{2f(m)}$ to ensure that $F(2x - m) \leq 1$. We can immediately recognize that our approximation is akin to the Uniform case, and conclude that $P(K_1 = k_1) \approx \frac{1}{2^{k_1+1}}$ for $k_1 < n-1$. Because this procedure applies to arbitrary $F(x)$, we can iterate this approximation for $K_2, K_3$ and so on. At each step, we will obtain a new distribution function by truncation, but the form of our approximation does not change (only the lower bound, $m'$, changes, and consequently $f(m')$ does, too).

This Taylor expansion argument provides an explanation for the wide applicability of our theory. By considering higher order terms in the Taylor expansion, we can also gain more general insight into deviations from our theory, such as those observed in the tails of the colonization rate distribution for the Exponential and Triangular distributions, above. Retaining the quadratic term in our approximation for $F(x)$, we have $F(x) \approx f(m)(x-m) + \frac{f'(m)}{2}(x-m)^2$ (and consequently $f(x) \approx f(m) + f'(m)(x-m)$), where $f'(m)$ is the second derivative of $F(x)$ with respect to $x$. Now consider $P(K_1 = 0)$. We have

$$P(K_1 = 0) \approx n \int_m^U \left(1 - 2f(m)(x-m) - 2f'(m)(x-m)^2\right)^{n-1} \left(f(m) + f'(m)(x-m)\right) dx\,.$$

As above, we define $U$ such that the integrand remains non-negative. Now observe that we can re-write this probability as

$$\begin{aligned} P(K_1 = 0) &\approx n \int_m^U \left(1 - 2f(m)(x-m) - 2f'(m)(x-m)^2\right)^{n-1} \left(f(m) + 2f'(m)(x-m)\right) dx\,, \\ &\quad - n \int_m^U \left(1 - 2f(m)(x-m) - 2f'(m)(x-m)^2\right)^{n-1} f'(m)(x-m)dx\,. \end{aligned} \tag{39}$$

Using the change of variables $y = 2f(m)(x-m) - 2f'(m)(x-m)^2$, it is easy to see that the first term is equal to $\frac{1}{2}$. Thus, the second term describes deviations from the expectation $P(K_1 = 0) = \frac{1}{2}$. The sign of this deviation is controlled by $f'(m)$. For $f'(m) > 0$, we expect $P(K_1 = 0) = \frac{1}{2}$ to overestimate the true probability. For $f'(m) < 0$, we expect the opposite. This is consistent with the deviations we have observed above: For the Triangular distribution, $f'(m) > 0$, and for the Exponential distribution, $f(x)$ is decreasing in the tail of the distribution. In fact, Eq. 39 can be evaluated exactly in cases where $f(m) = 0$ – such as the Triangular distribution – and we find that $P(K_1 = 0) = \frac{1}{2} - \frac{1}{4} = \frac{1}{4}$, as derived above.

Consideration of these Taylor approximations helps clarify when we will observe excess coexistence or exclusion. The linear approximation, developed above, will be poor when the term $f(m)(x - m)$ does not dominate higher order terms. This occurs when $f(m)$ is very small, or when $x$ is not very small. The latter can occur when $n$ is small, as in the right tail of the colonization rate distribution. When the linear approximation deteriorates, the sign of $f'(m)$ determines whether we will observe more or less competitive exclusion than expected. In regions of low but increasing probability density (i.e. $f(m) \ll 1$ and $f'(m) > 0$) there will be excess exclusion, while in regions of low, decreasing density (i.e. $f(m) \ll 1$ and $f'(m) < 0$) there will be excess coexistence. Both deviations can occur at different points for the same colonization rate distribution, as we show in Fig. S6 for truncated normal distributions. In such cases, these deviations will cancel out to some extent.

Additionally, while this approximation theory is based on the assumption of a continuous distribution, Fig. S7 shows that similar results hold even for more complex (mixture) distributions with a discontinuity.

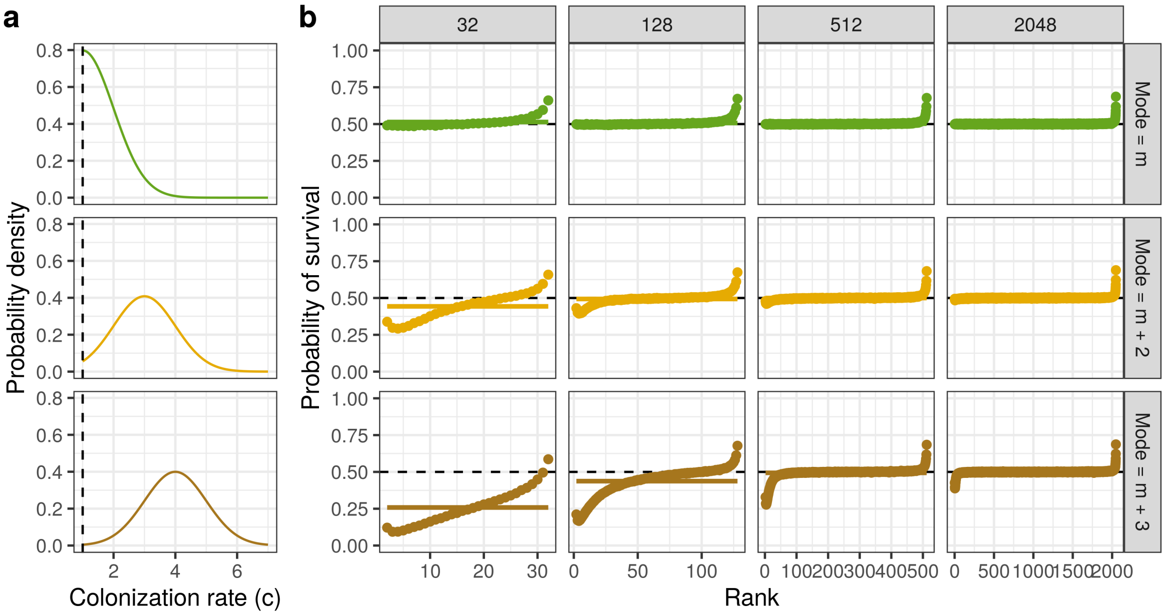


Figure S6: Probability of persisting in the assembled metacommunity by competitive rank in the pool (i.e. species 1 is the best competitor) for truncated normal distributions. (a) Probability density function for three different (left) truncated normal distributions, each with $\sigma = 1$ and tunctated at $m = 1$ (dashed line). The mode of the distribution is shifted in each case. (b) As for Fig. S5, but with colonization rates sampled from the truncated normal distributions shown in (a). Notice excess exclusion (coexistence) in the lower (upper) tails, as predicted by theory.

### S3 Spacing between colonization rates in the assembled metacommunity

In this section, we consider the distribution of gaps between colonization rates in assembled communities (that is, the quantities $c_i - c_{i-1}$ in the final metacommunity). Our linear niche shadow approximation (Eq. 31) implies that $\ell_i - c_i = c_i - \ell_{i-1} = X_i$. Thus, we have that the $c_i - c_{i-1} = c_i - \ell_{i-1} + \ell_{i-1} - c_{i-1} = X_i + X_{i-1}$. In Section S2.1, we developed the approximation that $X_i \sim \text{Exp}(nf(\ell_{i-1}))$. Using the (conditional) independence of $X_i$ and $X_{i-1}$, their sum is a two-parameter hypoexponential random variable with parameters $\lambda_1 = nf(\ell_{i-1})$ and $\lambda_2 = nf(\ell_{i-2})$. This distribution has the PDF

$$f_{\text{hypo}}(x) = \frac{\lambda_1 \lambda_2}{\lambda_1 - \lambda_2} \left( e^{-\lambda_2 x} - e^{-\lambda_1 x} \right) . \tag{40}$$

For the special case of colonization rates with a Uniform distribution on $(m, m+1)$, we have

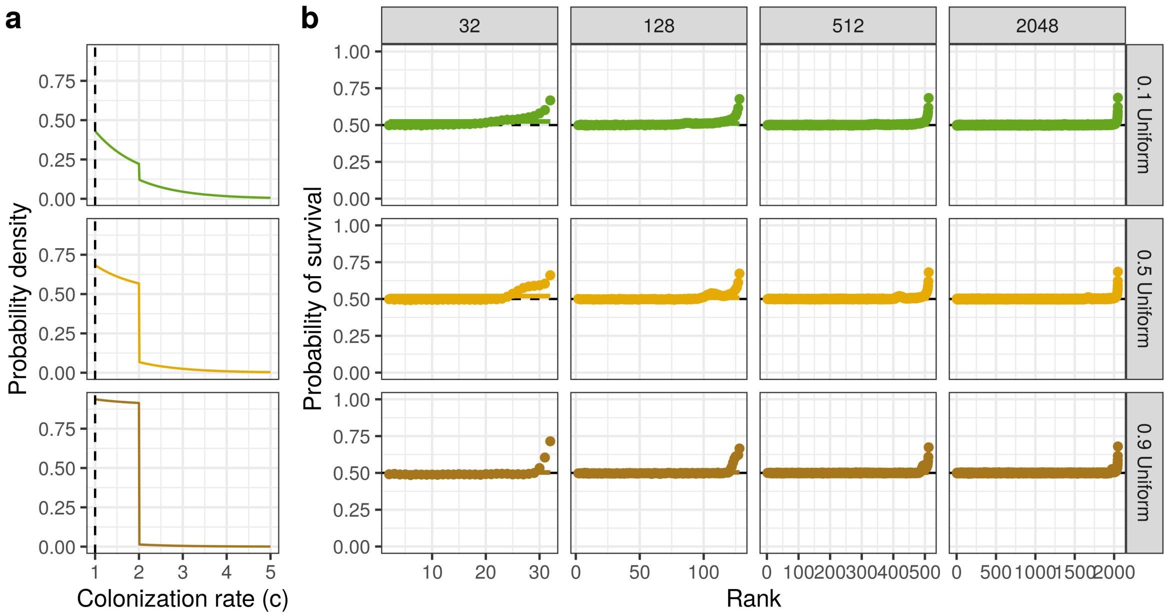


Figure S7: Probability of persisting in the assembled metacommunity by competitive rank in the pool (i.e. species 1 is the best competitor) for uniform-exponential mixture distributions. (a) Probability density function for three different mixutres of the uniform and exponential distributions, with the probability of sampling from the uniform distribution indicated. (b) As for Fig. S5, but with colonization rates sampled from the mixture distributions shown in (a). Although these distributions contain a discontinuity, results are qualitatively the same as our other cases.

$f(\ell_{i-1}) = f(\ell_{i-2}) = 1$ for all $i$. Under these conditions, $X_i + X_{i-1}$ has an Erlang distribution with shape parameter 2 and rate $\lambda = n$. As a point of comparison, we can compute the distribution of gaps that would be expected if species in the pool were chosen *independently* to persist in the final metacommunity. Conditioning on an observed final richness $s$, the colonization rates of the persisting species would be distributed as $s$ Uniform random samples. The gap between consecutive random samples in this case is distributed as $\text{Beta}(1, s)$[10]. As shown in Fig. S8, this distribution is qualitatively different from the Erlang distribution derived above. In particular, the Erlang distribution with shape parameter 2 has no mass at zero, while the Beta distribution $\text{Beta}(1, s)$ has its mode at zero. We observe that the Erlang prediction matches numerical simulations extremely well, while the Beta provides a poor match. This comparison provides a clear indication that the set of persisting species is overdispersed, relative to independent random sampling from the pool. As we discuss in the Main Text, the actual distribution of gaps is much more similar to the distribution of gaps between *every other* colonization rate in the pool. For uniformly distributed colonization rates, this distribution is $\text{Beta}(2, n-1)$. In the limit $n \to \infty$, $\text{Beta}(2, n-1) = \text{Gamma}(2, n-1)$, which is the Erlang distribution with shape parameter 2 and rate $n-1$. This distribution is therefore extremely close to the distribution derived for $c_i - c_{i-1}$ above.

More generally, we observe that Eq. 40, which holds for any distribution of colonization rates in the pool, has no mass at zero. Meanwhile, arguments similar to those in Section S2.1 can be used to establish that – for any distribution of colonization rates – the distribution of gaps between $s$ randomly selected species from a pool is approximately $\text{Exp}(sf(c_{i-1}))$ for each $c_i - c_{i-1}$. The Exponential distribution always has its mode at zero. Thus, we see that overdispersion in the persisting colonization rates is the typical behavior for any random pool.

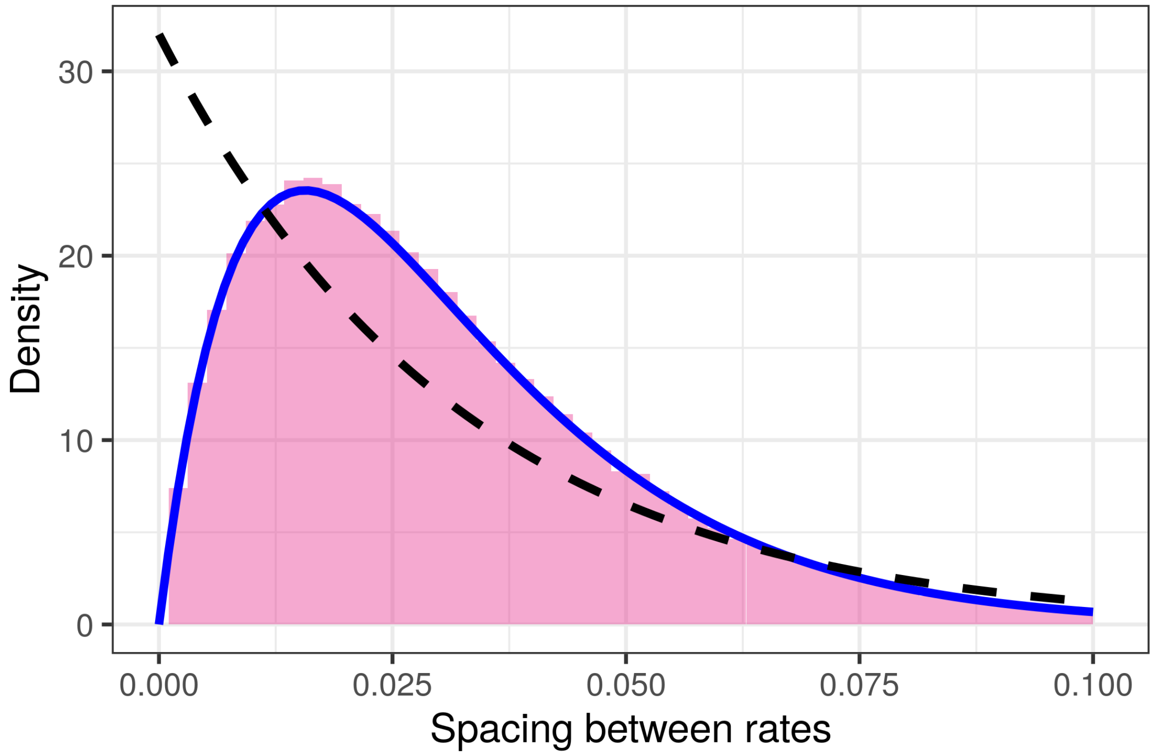


Figure S8: Distribution of gaps $c_i - c_{i-1}$ for Uniformly distributed colonization rates. Histogram shows the empirical distribution of spacings from $10^4$ assembled communities (resulting from random pools of $n = 64$ species). The theoretical distribution of spacings (Erlang distribution with shape parameter 2) is shown in blue. This prediction matches the simulation results extremely well. A naive prediction, Beta$(1, \frac{n}{2})$, corresponding to independent random sampling of $\frac{n}{2}$ species from the pool, is shown with black dashed line. This distribution has high density for small spacings, while the observed and theoretical distributions have very low density for small spacings, indicating overdispersion in the assembled colonization rates.

### S4 One-at-a-time assembly scenarios

#### S4.1 Fixed regional pool

In the all-at-once scenario, the dynamics of a pool of $n$ species is given by the differential equations of Eq. 1. The behavior is simple; independently of the parameters $(c, m)$ of the model and the initial condition, the dynamics converges to a unique equilibrium $p^\star = (p_1^\star, \ldots, p_n^\star)$. Direct information on the persistent species is given by the conditions of Eq. 25. We showed in Section S1.2 that there exists a unique globally stable equilibrium, i.e. if one waits long enough, the unique equilibrium is reached.

This equilibrium is called saturated because it is resistant against invasion of absent species. Let $i$ belong to the indices of the extinct species, then

$$\left(\frac{1}{p_i}\frac{dp_i}{dt}\right)_{p_i \to 0^+} \leq 0\,.$$

However depending of the construction process, the dynamics of Eq. 1 can be understood in two equivalent directions. On the one hand, there is a primary ecosystem with a pool of $n$ species that have different initial occupancies. The ecosystem changes continuously according to the ODE of the model. On the other hand, the ecosystem is assumed to be initially empty and species try to invade it sequentially in a random order. This one-at-a-time scenario generates disturbances to the pool of persisting species. When a species tries to invade the system at a certain time $t$, it can cause the extinction of other species, but also the expansion of other species that were already present in the system. Indeed, the particularity is that a species that has invaded the system, even if it becomes extinct, it can invade again at any time.

*Remark* 3. A permanent extinction never occurs in the dynamics of the model. However, the vanishing components corresponding to the species going to extinction with $p_i^\star = 0$ converges toward zero i.e. $p_i(t) \xrightarrow[t\to\infty]{} 0$.

For this model, both types of construction have the same final behavior; the two dynamics converge towards the same equilibrium which is true for infinitely long periods. Because there are never true extinctions in the dynamics of this model, this fact is guaranteed once each species has invaded the community at least once. However, the two assembly scenarios will converge even if we allow the possibility of some species going extinct (e.g. we take the metacommunity to be at its equilibrium state after each invasion). To see this, it is sufficient to observe that the final metacommunity can be assembled by introducing species one-at-a-time in the order of their colonization rates. Given enough invasions, this invasion sequence will eventually occur, at which point the equilibrium is non-invasible by the excluded species. In practice, however, convergence is usually much faster. Additionally, we note that the equivalence to a GLV model, shown in Section 1, is also sufficient to establish the equivalence of these two assembly scenarios, using the results of [11].

#### S4.2 De novo invasion

Here we derive a lower bound for the expected richness after $\tau$ invasion attempts. First, it is necessary to consider the fraction of invasion attempts that succeed, on average. Under our assumptions (namely, the assumption that $f(c) = 0$ for $c < m$), the first invasion succeeds with probability 1. The second invasion succeeds with probability at least $\frac{1}{2}$, regardless of the distribution of colonization rates. To see this, note that if the colonization rate of the first species is $x$, then the second invasion succeeds if the colonization rate of the invader is less than $x$ or greater than $2x - m$. The probability of this event is $1 - F(x^2/m) + F(x)$, or if the distribution has an upper bound $b$ and $x > \sqrt{bm}$, then the probability is simply $F(x)$ (because the invader cannot exceed the niche shadow of the first species; it can only outcompete this species). Integrating over all values of $x$, weighted by $f(x)$, gives

$$
\begin{aligned}
\int_m^\infty f(x)\left(1 - F\left(\frac{x^2}{m}\right) + F(x)\right)dx &= 1 - \int_m^\infty f(x)F\left(\frac{x^2}{m}\right)dx + \frac{1}{2}\,, \\
&\geq 1 - \int_m^\infty f(x)dx + \frac{1}{2}\,, \\
&= \frac{1}{2}\,,
\end{aligned}
$$

in the unbounded case and

$$
\begin{aligned}
\int_m^{\sqrt{bm}} & f(x)\left(1 - F\left(\frac{x^2}{m}\right) + F(x)\right) dx + \int_{\sqrt{bm}}^b f(x)F(x)dx \\
&= F(\sqrt{bm}) - \int_m^{\sqrt{bm}} f(x)F\left(\frac{x^2}{m}\right) dx + \frac{1}{2}F(\sqrt{bm})^2 + \left(\frac{1}{2} - \frac{1}{2}F(\sqrt{bm})^2\right) , \\
&\geq \frac{1}{2} + F(\sqrt{bm}) - \int_m^{\sqrt{bm}} f(x)dx , \\
&= \frac{1}{2} .
\end{aligned}
$$

Continuing this calculation for the success of the third invasion and so on is complicated by the fact that each new invader may cause the extinction of earlier species. Instead, we can "look ahead" to a state where many species have invaded. Once a large number of species are present in the metacommunity, the approximations developed in Section S2.1 hold. In particular, we expect that $\ell_i - c_i \approx c_i - \ell_{i-1}$ for all species $i$. An invader with colonization rate in the niche shadow, $(c_i, \ell_i)$, will fail, while an invader in the interval $(\ell_{i-1}, c_i)$ will succeed. Because the colonization rate axis can be partitioned into many such intervals of equal length, each invasion succeeds with probability near $\frac{1}{2}$ when there are many species in the metacommunity.

Overall, this leads us to expect that early in the assembly process more than half of invasions succeed, but the probability of success should approach one half over time. In other words, the number of successful invasions after a large number of attempts, $\tau$, will be approximately $\frac{\tau}{2}$ on average. However, the richness at this point will typically be much less than $\frac{\tau}{2}$, because many successful invasions will result in the extinction of some resident species. There is a limit, though, to the number of extinctions each species can induce. The hierarchical structure of this model implies that an invader can only cause the extinction of inferior competitors. To derive a lower bound on the average rate at which diversity accumulates, we can assume that each invader causes the extinction of *all* inferior competitors.

Let us first consider the species with the lowest colonization rate among the $\tau$ potential invaders. Obviously, this species succeeds in invading and is not displaced. Thus, this species is part of the persisting metacommunity after $\tau$ invasions. Because invaders are sampled independently from the underlying colonization rate distribution, it is equally likely that this species is any of the first $\tau' \leq \tau$ successful invaders. Under our "worst case" assumption, we suppose that if this species is the $i$th successful invader, it excludes all of the $i-1$ preceding species. Now consider the $\tau' - i$ successful invaders that arrive after the best competitor. One must be the best competitor among these species, and, as before, we can conclude that this species is not displaced up to the $\tau$th invasion. Thus, this species is also part of the persisting metacommunity. We can iterate this argument until the best competitor among the late invaders is the final one. This yields a set of "definitely persisting species" that provides a lower bound for the number of actual persisting species.

This picture leads to a recursive formula for the mean number of definitely persisting species after $\tau'$ successful invasion, $\mu_{\tau'}$, obtained by averaging over the possible positions of the best competitor at each step. We have

$$
\mu_{\tau'} = 1 + \frac{1}{\tau'} \sum_{i=1}^{\tau'} \mu_{\tau'-i} = 1 + \frac{1}{\tau'} \sum_{i=0}^{\tau'-1} \mu_i ,
$$

with $\mu_0 = 0$. Now we multiply both sides of this equation by $\tau'$, obtaining

$$
\tau' \mu_{\tau'} = \tau' + \sum_{i=0}^{\tau'-1} \mu_i ,
$$

and thus also

$$(\tau' - 1)\mu_{\tau'-1} = (\tau' - 1) + \sum_{i=0}^{\tau'-2} \mu_i \,.$$

Subtracting the latter from the former leaves us with

$$\tau'(\mu_{\tau'} - \mu_{\tau'-1}) + \mu_{\tau'-1} = 1 + \mu_{\tau'-1} \,.$$

Finally, rearranging yields

$$\mu_{\tau'} = \frac{1}{\tau'} + \mu_{\tau'-1} = \sum_{i=0}^{\tau'} \frac{1}{i} \,.$$

From this equation, we see that $\mu_{\tau'}$ is the harmonic number $H_{\tau'} \approx \log(\tau') + \gamma + \frac{1}{2\tau'}$, where $\gamma \approx 0.577$ is the Euler-Mascheroni constant. It is easy to verify that $H_{\tau'} > \log(1 + \tau')$ for $\tau' > 0$. Using the fact that $\tau'$ is typically very close to $\frac{\tau}{2}$ after $\tau$ invasion attempts, we obtain the lower bound $\mathbb{E}[S \mid \tau \text{ invasion attempts}] \geq \log(1 + \frac{\tau}{2})$.

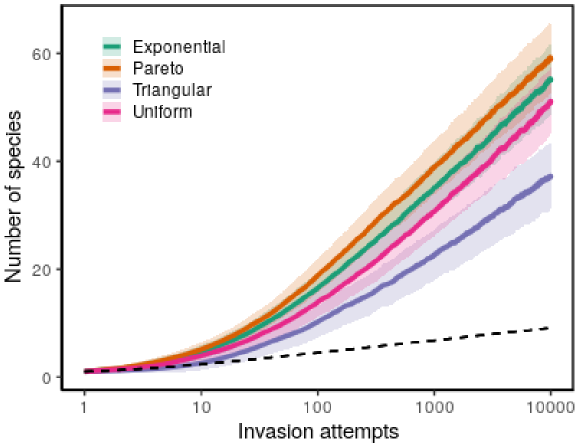


Figure S9: Accumulation of species richness in the de novo invasion scenario. As in Fig. 5b (Main Text), but for many more invasion attempts (note log scale for x-axis). Looking over very long assembly trajectories, the logarithmic growth of mean richness is apparent. The logarithmic lower bound derived in the text is shown as dashed black line. Curves represent statistical summaries of $10^3$ random assembly trajectories.

### S5 Equilibrium occupancies

In this section, we briefly consider the occupancy of each species in the equilibrium metacommunity. While our primary focus is on understanding the set of persisting species (*How many and which species persist?*), it is also natural to consider the composition of the metacommunity at equilibrium (*What proportion of patches are occupied, and what is the occupancy of each individual species?*).

As a first step, let us consider the total occupancy in the metacommunity, or equivalently, the proportion of patches unoccupied by any species at equilibrium. We denote by $h_i^\star$ the proportion of patches unoccupied by species $1 \ldots i$. Thus, the total occupancy at equilibrium is $1 - h_S^\star$. In a counter-intuitive way, even when the number of species is very large ($n \to \infty$), $h_S^\star$ does not necessarily converge towards 0. In the single-species Levins' model with parameters $(c, m)$, the fraction of empty patches is given by $m/c$. Thus, the higher the colonization rate, the smaller the proportion of empty patches (however, notice that this proportion is never zero unless $m = 0$ or $c \to \infty$). In the multispecies trade-off model, the conclusion is qualitatively similar; we have the relation $h_i^\star \approx (m/c_i)^{1/2}$ which can be derived as follows:

$$\begin{aligned} h_i^\star &= 1 - \sum_{j=1}^{i} p_j^\star\,, \\ &= \frac{m h_{i-1}^\star + \sum_{j=1}^{i-1} m p_j^\star}{c_i h_{i-1}^\star}\,, \\ &= \frac{m}{c_i h_{i-1}^\star}\,. \end{aligned}$$

Using the fact that $h_i^\star < h_{i-1}^\star$, one obtains the same inequality as in Kinzig *et al.*[12],

$$\sqrt{\frac{m}{c_{i+1}}} \leq h_i^\star \leq \sqrt{\frac{m}{c_i}}\,. \tag{41}$$

This equation shows that to have an idea of the empty space, it is sufficient to have a good approximation of $c_S$, which, for large $n$, will be close to the largest colonization rate in the pool, $c_n = \max_{i\in[n]} c_i$. For the cases we study (e.g. Uniform, Exponential, Pareto) the distributions of the maximum are generally known. The distributions with finite support are simplest to analyze. For any distribution with support on $(m, b)$ – assuming $f(x) > 0$ for all $m < x < b$ – if we consider $n$ *iid* random variables $U_i$, then $\mathbb{E}\left(\max_{i\in[n]} U_i\right) \xrightarrow[n\to\infty]{} b$.

When the distribution of the colonization rates $c$ has infinite support, this theory implies that when the number of species tends to infinity, the fraction of empty sites tends to zero. However, when $n$ is too small, the prediction is not accurate because of the gap between $c_{i+1}$ and $c_i$ (consider, for example, $n = 1$).

We can use the results above to study the distribution of occupancies of individual species. This distribution was considered by Kinzig *et al.* [12], who derived a relationship between occupancies and colonization rates: They showed that $p^\star(c) \propto c^{-3/2}$ when colonization rates are uniformly spaced along some interval. This power-law relationship implies that the most competitive species tend to occupy a larger share of the landscape than the best colonizers.

Using the results above, we can derive a more general relationship between occupancies and colonization rates which applies to our probabilistic setting. From Tilman [1], we have the equality

$$p_i^\star = 1 - \sum_{j=1}^{i-1} p_j^\star - \frac{m + \sum_{k=1}^{i-1} c_k p_k^\star}{c_i}\,.$$

Using our definition of $h_i^\star$ and re-arranging, we have

$$\begin{aligned} p_i^\star &= (h_{i-1}^\star)\left(1 - \frac{1}{c_i}\frac{m + \sum_{k=1}^{i-1} c_k p_k^\star}{h_{i-1}^\star}\right)\,, \\ &= h_{i-1}^\star \frac{1}{c_i}(c_i - \ell_{i-1})\,. \end{aligned}$$

This last inequality comes from recognizing a definition of the threshold value $\ell_{i-1}$, which is a direct consequence of Eq. 4 in Section 1. Earlier, we denoted the difference $c_i - \ell_{i-1}$ by $X_i$; using this notation we have

$$p_i^\star = \frac{h_{i-1}^\star X_i}{c_i}\,.$$

Plugging in the approximation $h_i^\star \approx (m/c_i)^{1/2}$ recovers the power-law of Kinzig *et al.*, up to a factor of $X_i$. Intuitively, this dependence on $X_i$ tells us that species that are very similar to the next best colonizer ($c_i$ close to $\ell_{i-1}$) have lower occupancy than species that are more dissimilar. In the same way that the niche shadow cast by species $i$ depends on this gap, so does its occupancy. When a species is close to its threshold value, $\ell_{i-1}$, it has high niche overlap with the superior competitor, and it is competitively suppressed as a result.

For large assembled metacommunities ($n \to \infty$), we can develop a prediction for the distribution of occupancy, as a function of competitive rank. Specifically, we treat competitive rank as a continuous variable, using $x \approx i/S$. We expect that $c_i \approx F^{-1}(x)$, at least for distributions with bounded support. Combining this fact with the approximation we derived for $X_i$ in Section 2, $X_i \sim \mathrm{Exp}(nf(\ell_{i-1}))$, and additionally using $\ell_{i-1} \approx c_i$, we have an approximation for the distribution of occupancies at equilibrium, as a function of $x$:

$$p^\star(x) \sim \mathrm{Exp}\left(n\, f\left(F^{-1}(x)\right)\,\left(F^{-1}(x)\right)^{\frac{3}{2}}\right)\,. \tag{42}$$

For the Uniform distribution on $(m, m+1)$, for example, this reduces to

$$p^\star(x) \sim \mathrm{Exp}\left(n\,(x+m)^{\frac{3}{2}}\right)\,.$$

This random variable has the conditional mean $n^{-1}(x+m)^{-3/2} \approx n^{-1}c_i^{-3/2}$, as anticipated. In Figs. S10-S13, we plot the relationship between occupancies and competitive rank (in the pool) for all four of our example distributions.

It is interesting to compare Figs. S10-S13 with Fig. S5. While we have seen that all species in the pool persist with equal probability, here we find that they systematically differ in their equilibrium occupancies. The relationship between competitive rank (or colonization rates) and occupancy also differs depending on the distribution of colonization rates. For all distributions, the 3/2 power factor in Eq. 42 tends to increase the occupancy of better competitors. However, there is also an effect of the density, which we see clearly in Figs. S11-S13. In regions of the distribution with low density, the gaps $X_i$ tend to be larger, promoting higher occupancies.

### S6 Definition of standard distributions

For reproductibility purposes, we recall some standard positive probability distributions (see the handbook of Abramowitz [9]). Each distribution is defined by a random variable $X$ following the probability distribution function (PDF) $f$ and the cumulative distribution function (CDF) $F$. We denote by

$$\mathbf{1}_{[a,b]}(x) = \begin{cases} 1 \text{ if } x \in [a,b], \\ 0 \text{ else,} \end{cases}$$

the characteristic function.

**Definition 1** (Continuous Uniform)**.** The continuous Uniform distribution $\mathcal{U}([a,b])$ describes an experiment where there is an arbitrary outcome that lies between certain bounds: $(a,b) \in \mathbb{R}_+^2$,

$$f(x;a,b) = \frac{1}{b-a}\mathbf{1}_{[a,b]}(x)\ ,\ \ F(x;a,b) = \frac{x-a}{b-a}\mathbf{1}_{[a,b]}(x) + \mathbf{1}_{[b,+\infty)}(x).$$

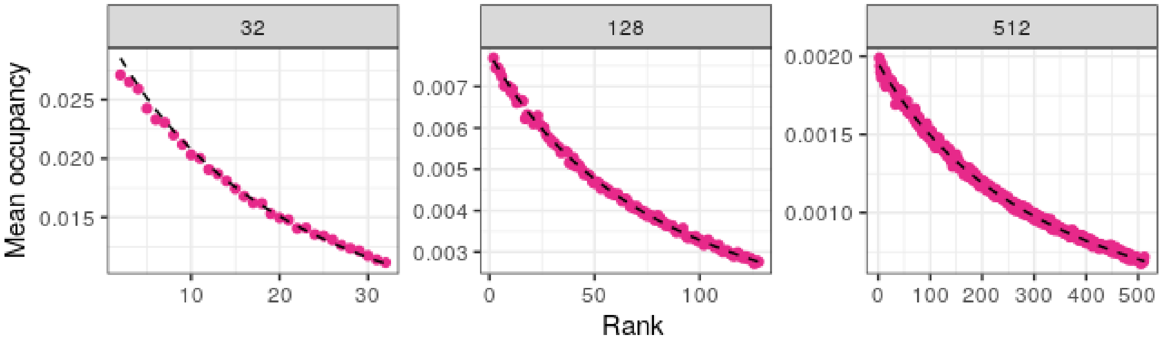


Figure S10: Mean occupancy (conditional on presence) in the assembled metacommunity by competitive rank for uniformly distributed colonization rates. We show results for 3 different pool sizes (values of $n$), representing averages over $10^4$ random pools. The theoretical prediction, given by the conditional mean of Eq. 42, is indicated by a dashed black line.

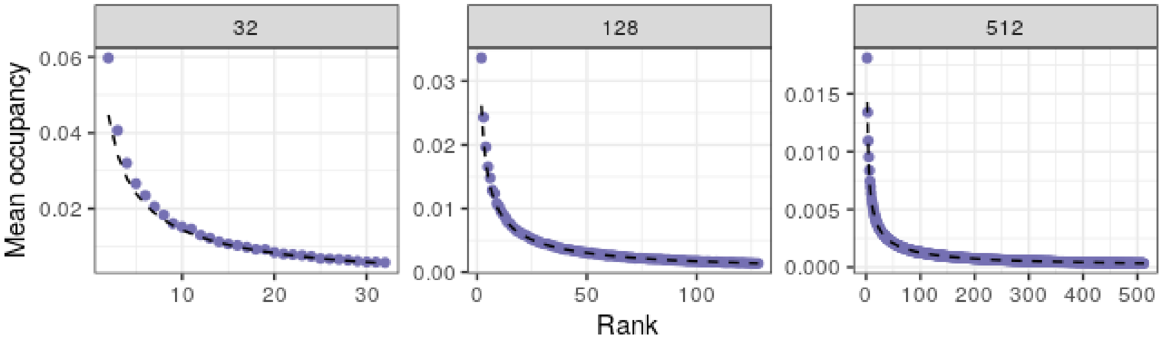


Figure S11: As in Fig. 42, but for the Triangular distribution.

Given $X$ a random variable following the distribution $\mathcal{U}([a, b])$

$$\mathbb{E}(X) = \frac{a+b}{2} \ , \ \mathrm{Var}(X) = \frac{(b-a)^2}{12} \, .$$

Throughout this study, we take $a = m$ unless otherwise specified. In all simulations and figures, we additionally use $b = m + 1$.

**Definition 2** (Pareto)**.** The Pareto distribution $\mathcal{P}(a)$ is a power-law distribution with shape $a$, support $[1, +\infty)$,

$$f(x; a) = \frac{a}{x^{a+1}} \mathbf{1}_{[1,+\infty)}(x) \ , \ F(x; a) = \left(1 - \frac{1}{x^a}\right) \mathbf{1}_{[1,+\infty)}(x).$$

Given $X$ a random variable following the distribution $\mathcal{P}(a)$:

$$\mathbb{E}(X) = \begin{cases} \frac{a}{a-1} \text{ if } a > 1, \\ +\infty \text{ otherwise} \end{cases} , \ \mathrm{Var}(X) = \begin{cases} \frac{a}{(a-1)^2(a-2)} \text{ if } a > 2, \\ +\infty \text{ otherwise.} \end{cases}$$

The Pareto PDF $f_{par}$ can be expressed as a function of the Exponential PDF $f_{\exp}$:

$$f_{par}(x; a) = f_{\exp}(\log(x); a) \, .$$

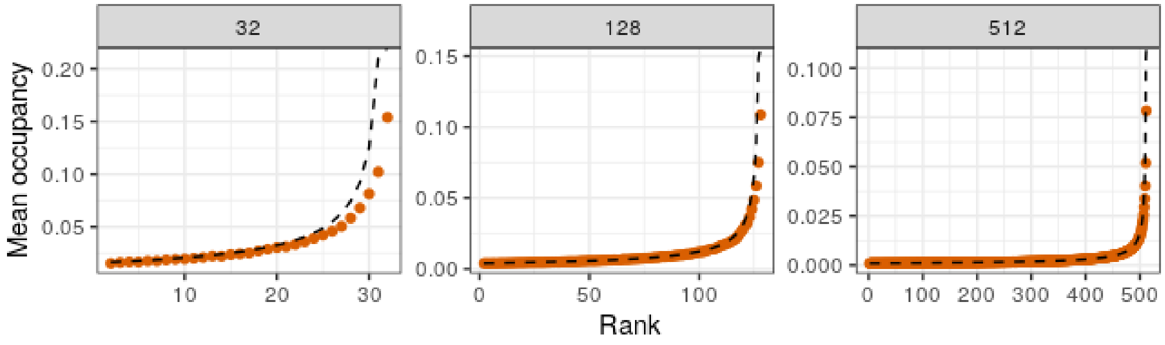


Figure S12: As in Fig. 42, but for the Pareto distribution.

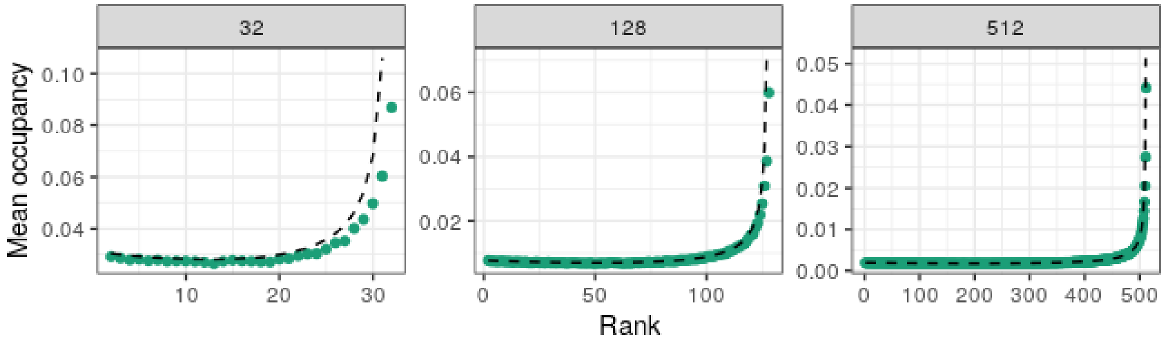


Figure S13: As in Fig. 42, but for the Exponential distribution.

**Definition 3** (Triangular)**.** The Triangular distribution $\mathcal{T}(a, b)$ is a probability distribution with a lower limit $a$, an upper limit $b$ whose density form a triangle:

$$f(x; a, b) = \frac{2(x-a)}{(b-a)^2}\mathbf{1}_{[a,b]}(x) \ , \ F(x; a, b) = \frac{(x-a)^2}{(b-a)^2}\mathbf{1}_{[a,b]}(x) + \mathbf{1}_{[b,+\infty)}(x).$$

Given $X$ a random variable following the Triangular distribution $\mathcal{T}(a, b)$:

$$\mathbb{E}(X) = \frac{a+2b}{3} \ , \ \mathrm{Var}(X) = \frac{a^2+b^2-2ab}{18} \, .$$

Throughout this study, we take $a = m$ unless otherwise specified. In all simulations and figures, we additionally use $b = m + 1$.

**Definition 4** (Exponential)**.** The Exponential distribution $\mathrm{Exp}(\lambda)$, $\lambda > 0$ is a continuous analogue of the geometric distribution.

$$f(x, \lambda) = \lambda e^{-\lambda x}\mathbf{1}_{[0,+\infty)}(x) \ , \ F(x, \lambda) = 1 - e^{-\lambda x}\mathbf{1}_{[0,+\infty)}(x).$$

Given $X$ a random variable following the distribution $\mathrm{Exp}(\lambda)$:

$$\mathbb{E}(X) = \frac{1}{\lambda} \ , \ \mathrm{Var}(X) = \frac{1}{\lambda^2} \, .$$

Throughout this study, when we refer to exponentially-distributed colonization rates, we refer to a shifted (equivalently, truncated) version of the Exponential distribution, such that $c\,X + m$, where $X$ is an Exponential random variable. Additionally, in all simulations and figures, we use $\lambda = 1$.

[3] Lotka, Alfred. *Elements of Physical Biology.* Williams and Wilkins Company, 1925.

[9] Milton Abramowitz. *Handbook of mathematical functions: with formulas, graphs, and mathematical tables.* U. S. Government Printing Office, 1972.

[10] Herbert A David and Haikady N Nagaraja. *Order statistics.* John Wiley & Sons, 2004.

[11] Carlos A. Serván and Stefano Allesina. Tractable models of ecological assembly. *Ecology Letters*, 24(5):1029–1037, 2021.

[12] A. P. Kinzig, S. A. Levin, J. Dushoff, and S. Pacala. Limiting similarity, species packing, and system stability for hierarchical competition-colonization models. *The American Naturalist*, 153(4):371–383, 1999.
